# Capturing non-local through-bond effects in molecular mechanics force fields I: Fragmenting molecules for quantum chemical torsion scans [Article v1.1]

**DOI:** 10.1101/2020.08.27.270934

**Authors:** Chaya D Stern, Christopher I Bayly, Daniel G A Smith, Josh Fass, Lee-Ping Wang, David L Mobley, John D Chodera

**Author notes:** **For correspondence:** (CDS); (JDC).

## Abstract

Accurate molecular mechanics force fields for small molecules are essential for predicting protein-ligand binding affinities in drug discovery and understanding the biophysics of biomolecular systems. Torsion potentials derived from quantum chemical (QC) calculations are critical for determining the conformational distributions of small molecules, but are computationally expensive and scale poorly with molecular size. To reduce computational cost and avoid the complications of distal through-space intramolecular interactions, molecules are generally fragmented into smaller entities to carry out QC torsion scans. However, torsion potentials, particularly for conjugated bonds, can be strongly affected by through-bond chemistry distal to the torsion itself. Poor fragmentation schemes have the potential to significantly disrupt electronic properties in the region around the torsion by removing important, distal chemistries, leading to poor representation of the parent molecule’s chemical environment and the resulting torsion energy profile. Here we show that a rapidly computable quantity, the fractional Wiberg bond order (WBO), is a sensitive reporter on whether the chemical environment around a torsion has been disrupted. We show that the WBO can be used as a surrogate to assess the robustness of fragmentation schemes and identify conjugated bond sets. We use this concept to construct a validation set by exhaustively fragmenting a set of druglike organic molecules and examine their corresponding WBO distributions derived from accessible conformations that can be used to evaluate fragmentation schemes. To illustrate the utility of the WBO in assessing fragmentation schemes that preserve the chemical environment, we propose a new fragmentation scheme that uses rapidly-computable AM1 WBOs, which are available essentially for free as part of standard AM1-BCC partial charge assignment. This approach can simultaneously maximize the chemical equivalency of the fragment and the substructure in the larger molecule while minimizing fragment size to accelerate QC torsion potential computation for small molecules and reducing undesired through-space steric interactions.

## 1 Introduction

Molecular mechanics (MM) small molecule force fields are essential to molecular design for chemical biology and drug discovery, as well as the use of molecular simulation to understand the behavior of biomolecular systems. However, small molecule force fields have lagged behind protein force fields given the larger chemical space these force fields must cover to provide good accuracy for druglike ligands, common metabolites, and other small biomolecules [18, 52]. Torsion potentials for small molecule force fields are particularly problematic because historical approaches to their determination tend to produce parameters that generalize poorly [45]. In particular, torsion potentials where the central bond is part of a conjugated system of bonds can be strongly influenced by distal substituents, an effect very difficult to represent in a force field using only local chemistry to define the parameters. This lack of generalizability has led many practitioners to aim to improve the force field accuracy by refitting torsion parameters for individual molecules in a semi-automated bespoke fashion [4, 21, 31]. The added cost and time for this leads to a significant barrier to setting up simulations for new projects and may not produce generalizable parameters.

In many molecular mechanics force fields (such as Amber [7], CHARMM [5], OPLS [22]) a low-order Fourier series, such as a cosine series, is used to represent the contribution of torsion terms to the potential energy:

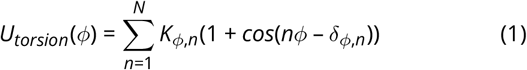

Here, *K*_*ϕ,n*_ is the torsion force constant which determines the amplitudes, *n* is the multiplicity which determines the number of minima, and *δ* is the phase angle which is sometimes set to 0*°* or 180*°* to enforce symmetry around zero. In most force fields, the maximum periodicity included is *N* = 4, though some force fields feature up to *N* = 6. Torsion potential parameters, such as the amplitudes *K*_*ϕ,n*_ and phase angle *δ*, are generally fit to the residual difference between a QC torsion energy profile (in the gas phase or implicit solvent) and the potential created by the non-torsion MM parameters [17]. The QC torsion energy profile is generated by fixing the torsion angle and geometry optimizing all other atomic positions.

This simple Fourier expansion of a single torsion angle is only an approximation to the true Born-Oppenheimer surface; neighboring torsions can have correlated conformational preferences the low-order Fourier series does not capture [9]. To account for this, 2D spline fits, such as the CMAP potential [29, 44], have become a popular way to model non-local correlations by fitting residuals between a 2D QC torsion energy profile of the coupled torsions of interest and the 2D MM torsion energy profile.

To produce a quantum chemical energy profile representing the chemical environment around the torsion, molecules are generally fragmented to smaller, model fragments and then capped with hydrogen or methyl groups for two main reasons described below.

### 1. Computational efforts scale poorly with molecule size

Generating 1D QC torsion profiles is computationally expensive, and becomes increasingly costly for larger molecules and/or higher dimensional QC torsion profiles. QC calculations scale poorly with the number of basis sets *N*, as *O*(*N*^*M*^). Formally, *M* ≤ 4 for hybrid DFT, although modern implementations of hybrid DFT scale asymptotically as *N*^2.2–2.3^ [43]. Using QCArchive data [39], we found that empirically, hybrid DFT grows like *N*^2.6^ for the DZVP basis [10] as shown for gradient evaluations in ***Figure 1***A. To achieve good sampling of the torsion profiles for adequate potential fits, constrained geometry optimizations need to be calculated at ≤ 15^0^ intervals, for a minimum of 24 constrained geometry optimizations for a 1D torsion profile^1^. To avoid hysteresis in the energy profile due to orthogonal degrees of freedom [46], methods like wavefront propagation [37] are used. This adds a factor of 2 ∗ *D*, where *D* is the dimension of the torsion scan, to the number of required optimizations. We found (SI ***Figure 4***) that on average, 60 QC optimizations are needed for a 1D wavefront-propagated torsion drive to converge, with 20 gradient evaluations required per optimization (SI ***Figure 3***). ***Figure 1***B shows a smoothed histogram of the distribution of the number of heavy atoms in FDA approved small molecules taken from DrugBank [51]. For an average druglike molecule of 25 heavy atoms (***Figure 1***B), we can estimate the cost of a 1D torsion scan to be 60 ∗ 20 ∗ 0.26 ∗ 25^2.6^ ≈ 1, 000, 000 CPU seconds. Reducing the size of the molecule to be used in QC torsion drive computations to 15 heavy atoms would reduce the cost the torsion scan by an order of magnitude (60 ∗ 20 ∗ 0.26 ∗ 15^2.6^ ≈ 300, 000 CPU seconds)

**Figure 1.**
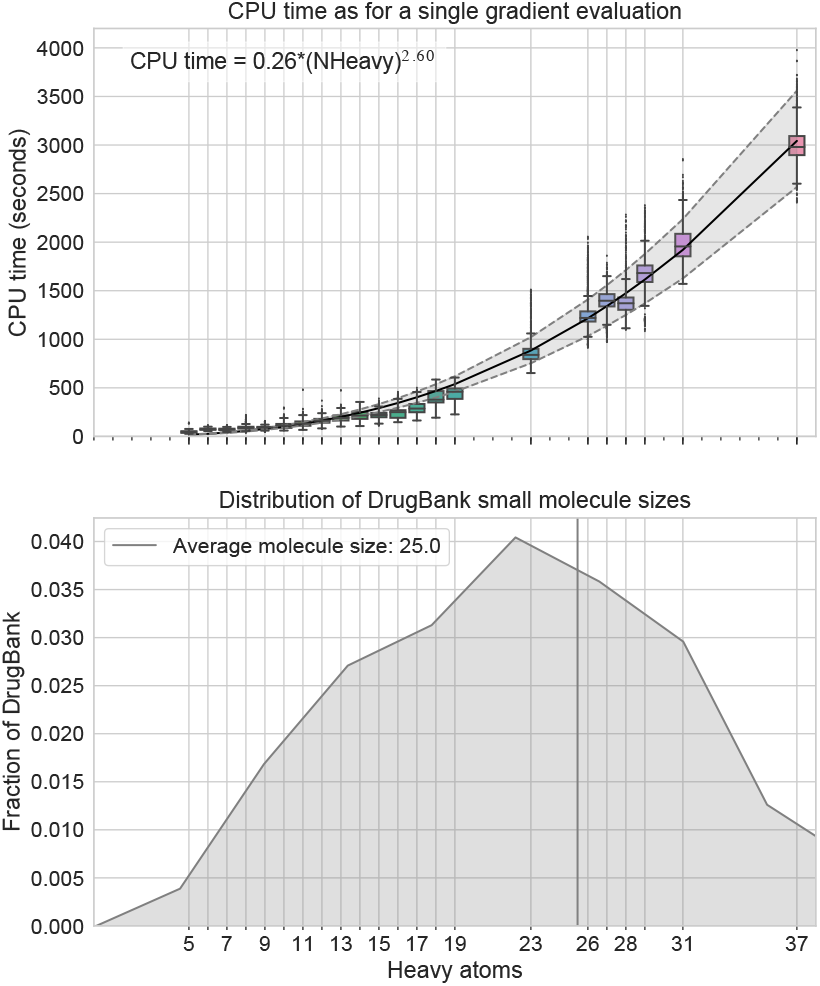
Fragmenting molecules is necessary to avoid high computational cost of generating QC data. **[A]** CPU time (wall clock time * nthreads) for a single QC gradient evaluation at B3LYP-D3(BJ)/DZVP level of theory [10, 15] vs number of heavy atoms in molecules. All computations shown here were run on an Intel(R) Xeon(R) CPU E5-2697 v4 @ 2.30GHz. Empirically, gradient evaluations grow as *O*(*N*^2.6^) where *N* is the number of heavy atoms. The scaling is similar on other processors shown in SI ***Figure 1 &*** SI ***Figure 2***. The black curves shows a power law fit to the data, while the grey curve depicts the 95% CI of the curve estimate. **[B]** Smoothed histogram of heavy atoms in small molecules from DrugBank (*≈* 10, 000 small molecules). The average small molecule in DrugBank has 25 heavy atoms as shown with the grey line.

### 2. Intramolecular interactions complicate torsion drives and torsion parameter fitting

In larger molecules, there is a greater potential for long-range through-space intermolecular interaction (e.g hydrogen bonding) that should be captured by non-bonded parameters, to contribute to the torsion profile. This can be difficult to decouple from the 1-4 interactions that torsion parameters are supposed to capture. While this can also happen in smaller molecules such as ethylene glycol [37] this problem is reduced when a minimal model molecule containing the model chemistry around the torsion is used, albeit not eliminated.

While several algorithms for fragmenting large molecules into smaller molecular fragments have been previously proposed, few are appropriate for generating high-quality torsion scans. Many of these algorithms fall into two categories:

1. fragmenting molecules for synthetic accessibility [3, 25, 26]
2. fragmenting molecules to achieve linear scaling for QC calculations [6, 8, 13, 35]

Fragmentation schemes for synthetic accessibility find building blocks for combinatorial and fragment-based drug design. Cleavage happens at bonds where it makes sense for chemical reactions and does not consider how those cuts affect the electronic properties of the fragments. For retrosynthetic applications, many cleavage points are at functional groups because those are the reactive sites of molecules. However, for our application, we especially do not want to fragment at these reactive points given their likelihood of having strong electron-donating or withdrawing effects which in turn strongly impact the torsional potential targeted for parameterization. Fragmentation algorithms for linear-scaling quantum chemical approaches such as Divide-and-Conquer methods [6], effective fragment potential method [24], and systematic molecular fragmentation methods [33] require the users to manually specify where the cuts should be or which bonds not to fragment. Furthermore, besides the scheme suggested by Rai et al. [38], none of these methods address the needs specific to fragmenting molecules for QC torsion scans for fitting MM force field parameters. When fragmenting molecules for QC torsion scans, fragments need to include all atoms involved in 1-4 interactions, since they are incorporated in the fitting procedure. We also need a systematic way to determine if remote substituents significantly alter the barrier to rotation around the central torsion bond.

In this work, we use the Wiberg Bond Order (WBO) [49], which is both simple to calculate from semi-empirical QC methods and is sensitive to the chemical environment around a bond [42]. WBOs quantify electron population overlap across bonds or the degree of bonding between atoms. Bond orders are correlated with bond vibrational frequencies [28, 50] and WBOs are used to predict trigger bonds in high energy-density material because they are correlated with the strength of bonds [19], a property which also directly affects the torsion potential. Wei et al. [48] have shown that simple rules for electron richness of aromatic systems in biaryls are a good indication of torsion force constants (specifically for *K*_*ϕ,n*_ in equation 1); however, this measure was only developed for biaryls and it does not take into account substituents beyond the aromatic ring directly adjacent to the bond.

Here, we develop an approach that uses the WBO to validate whether a fragmentation scheme significantly alters the local chemical environment of interest, with a particular focus on fragmentation schemes suitable for QC torsion drives. Our approach uses simple heuristics to arrive at a minimal fragment for QC torsion scan that is representative of the torsion environment of the substructure in the parent molecule: For a central bond, all atoms that are involved in the local steric interactions of a torsion scan are included, and then the WBO is used as a surrogate to determine if the fragment needs to be expanded to preserve the correct electronics around the central bonds.

The paper is organized as follows: Section 2 provides a mathematical and physical description of the problem of fragmenting molecules for QC torsion scans. Section 3 provides the motivation for using the WBO as a surrogate, evaluates its robustness, proposes a minimal fragmentation scheme, and describes a rich validation set that can be used to benchmark fragmentation schemes. Section 4 provides a discussion of the implications of this study, and Section 5 describes the detailed methods used here.

## 2 Theory and definitions

### 2.1 A mathematical definition of the problem of fragmenting molecules for QC torsion drives

A molecular structure can be modeled as a degree-bounded graph *G* = (*V, E*), where *V* are the nodes and *E* are the edges. In a molecular graph, the nodes correspond to atoms and edges correspond to covalent bonds. We define rotatable bonds as a set of bonds in the molecule that are not in rings, ℰ ⊂ *E*, and 𝒢 as a set of allowable subgraphs, where each subgraph *G*′ (*V*′, *E*′) ∈ 𝒢 is built around a central, rotatable bond, *e*′ ∈ ℰ with the following conditions:

1. The number of atoms in *G*′, |*V*′|, are 4 ≤ |*V*′| ≤ |*V* |
2. The minimum edges in *G*′ are *e*′ and all *e* ∈ *E* such that *e, e*′ share a vertex
3. All *v* ∈ *V* adjacent to *e* are included in *G*′
4. If *v* ∈ *V*′ is in a ring in *G*, the ring should be included in *G*′

The weights on *e*′ are given by *δ*(*e*′; *G′*), where *δ* is the RMSE of the torsion potential around the central, rotatable bond in the full graph *G* compared to the subgraph *G′*. Since *δ*(*e′*; *G′*) is computationally expensive to evaluate, we use a surrogate, *γ*(*e′*; *G*′), which we define as the difference of the WBO on the central, rotatable bond *e*′ in *G*′ and in the full graph *G*. In order to calculate the WBO, we need to “cap” open valences by adding additional atoms to ensure the resulting molecule is not a radical. The rules we use are defined in Section 3.2.

We want to minimize *γ*(*e*′; *G*′), while also minimizing the cost of each subgraph. We define the cost as estimated in ***Figure 1***.

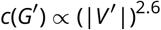

which leads to minimizing

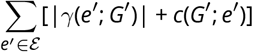

The search space of 𝒢 for each rotatable bond is combinatorial, and its upper bound is 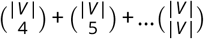 since all *G*′ ∈ 𝒢 need to be connected and rings are not fragmented. To reduce the search space, we also define a list of functional groups that should not be fragmented in Table 1.

**Table 1.**
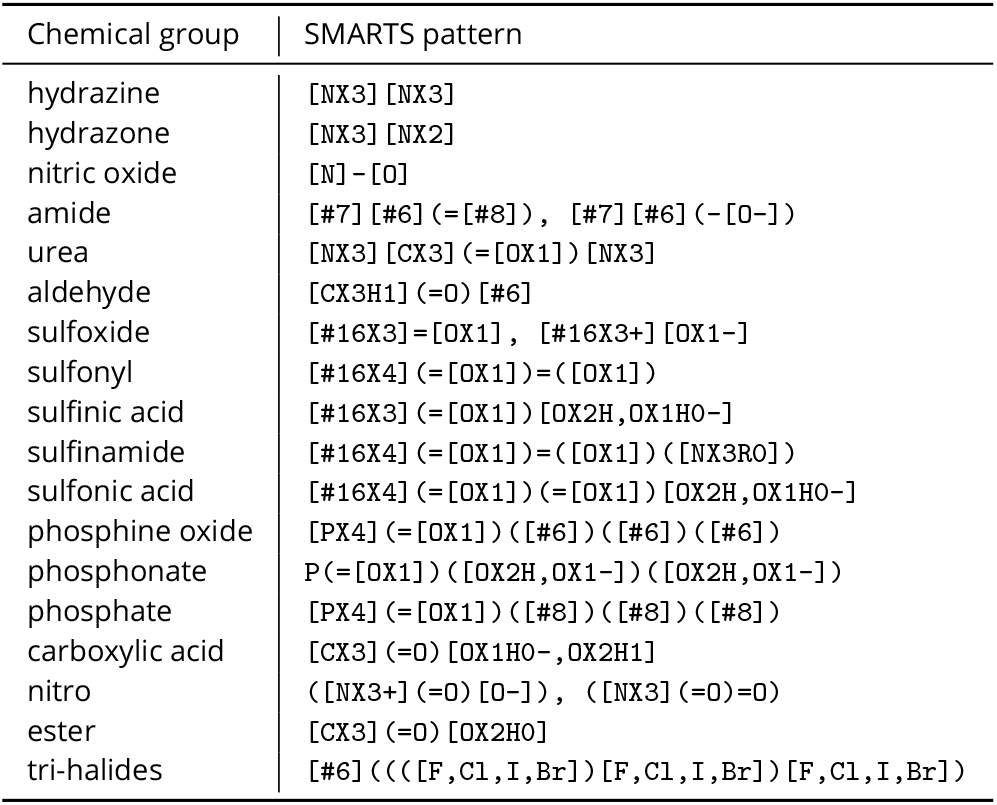
Functional groups kept whole during fragmentation. This list includes functional groups that were present in the validation set. Users can add their own functional groups they do not want to fragment.

Given how large the search space can become, we use several heuristics, described in Section 3.2.

### 2.2 Physical definitions

In most MM force fields, atom types are used to encode the local chemical environment of atoms for which parameters are assigned to [52]. This allows parameters developed for one atom to be transferred to another atom in a similar chemical environment. However, since atom types are locally defined, they do not always capture long-range effects such as conjugation. This is especially problematic for torsions since the torsion energy function (or profile) of a bond is determined by a combination of local and non-local effects such as conjugation, hyperconjugation, sterics, and electrostatics [12, 23, 27, 36, 48]. In this study, we define *local atoms* as those within two bonds of the central bond of a torsion, and *remote atoms* as any atom beyond those two bonds.

Steric and electrostatic interactions are, in principle, accounted for by non-bonded terms in most force fields, so a torsion profile would ideally primarily capture conjugation or hyperconjugation, and only the 1-4 nonbonded effects. Using small fragments to generate QC torsion profiles reduces non-local electrostatics and steric interactions from contributing to the torsion profile. However, conjugation and hyperconjugation are non-local properties and remote chemical changes can influence the extent of conjugation and/or hyperconjugation. In this study, we aim to avoid removing remote chemical substituents that impact the torsion of interest via conjugation and hyperconjugation by first understanding how the strength of the central bond is altered by remote chemical modifications. Below, we give a more precise definition of *conjugation* and *hyperconjugation* and how we use these terms in this paper.

Conjugation and hyperconjugation describe the sharing of electron density across several bonds. Conjugation is formally defined as the overlap of p-orbital electrons across *σ* bonds [Goldbook], such as what occurs in benzene or butadiene. Hyperconjugation is the interaction of electrons in a donating bonding orbital to an anti-bonding orbital [32]. In this study, for simplicity, we use the term conjugation to refer to all modes of conjugation and hyperconjugation.

## 3 Results and Discussion

### 3.1 Fragmenting molecules for quantum chemical torsion scans can alter the chemical environment of the torsion

Small chemical changes to molecules, both local and remote, can drastically alter their electronic properties. This poses a challenge to force field parametrization, specifically that of torsion parameters which rely on computationally expensive QC torsion scans for reference data [23]. Given how poorly QC torsion scans scale with molecular size, molecules are fragmented to smaller entities to generate the reference data. The assumption is that torsion scans generated for substructures can be used as reference data for other molecules that share the substructure. However, seemingly similar torsion types in different chemical environments can have very different torsion potential. This is particularly apparent in conjugated systems where small, remote changes can exert strong electron-donating or -withdrawing effects thus altering torsional barrier heights When such remote changes are combined with other more local changes that induce both electronic effects and steric clashes, remote effects can be either amplified or counteracted.

We illustrate such changes in an example as shown in ***Figure 2. Figure 2*** A shows a series of biphenyls in different protonation states. The central bond in the biphenyls are part of a larger conjugated system. However, the extent of conjugation changes with small, remote chemical changes. This is reflected both in the increased barrier heights of the torsion scans and increased Wiberg bond order of the bond. As the extent of conjugation increases, the Wiberg bond order, or electronic population overlap increases, and resistance to rotation increases as reflected in increasing torsion barrier heights. For ***Figure 2*** A, the remote change exerts a very strong effect on the central bond. This effect decreases with small local chemical changes that induce steric effects as shown in ***Figure 2*** B and C. When halogens, which are larger than hydrogens, are placed in the ortho position, the local steric effect increases the energy of the planar conformation needed for conjugation which counteracts the remote effects. Thus, while the energy barrier and WBOs increase for the cation, anion and zwitterion, the changes are less drastic for larger substituents at the ortho position. ***Figure 2*** C shows the series of torsion scans for chloro-pyridyl-phenol. While the torsion barrier heights increase as the WBO increase, the increase is less drastic as the flouro-pyridyl-phenol series (***Figure 2*** B) because chlorine is larger than fluorine.

**Figure 2.**
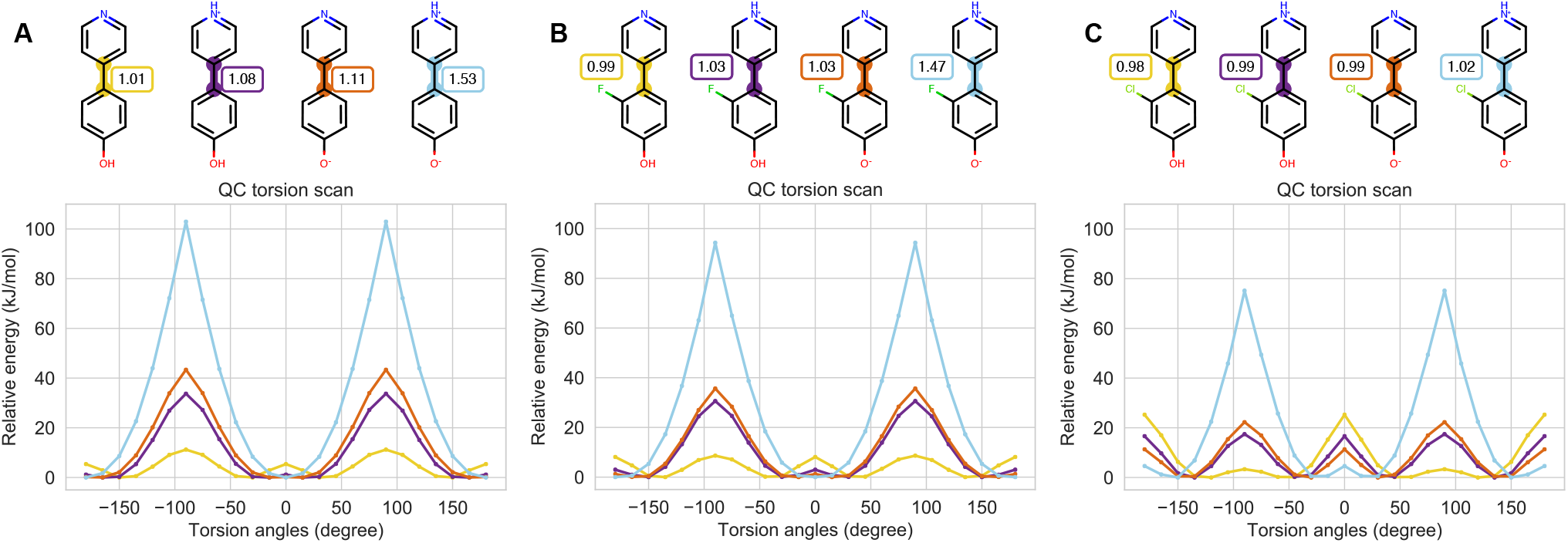
Torsion profiles can be sensitive to small local and remote chemical changes. **[A]** QC torsion scans of the central bond in biphenyls in different protonation states. The colors of the QC torsion scans correspond to the highlighted central bond of the biphenyls. The Wiberg bond order is shown in the box next to the central bond. **[B]** Same as **A** but with fluorine on the ortho position. **[C]** Same asA but with chlorine on the ortho position

The trends shown in this example illustrate why fragmenting molecules appropriately for QC torsion scans currently still requires human expertise and is difficult to automate. In these cases, small remote changes gradually perturbed the conjugation of the central bond. In addition, these changes were different for similar molecules with different local environments. When fragmenting molecules, we aim to avoid destroying a bond’s chemical environment by naively removing an important remote substituent.

### 3.2 A simple fragmentation scheme can use the WBO to preserve the chemical environment around a torsion

The WBO is a robust indicator of changes in torsion energy barrier heights for related torsions Therefore, if a fragment’s WBO deviates too much from its parent WBO at the same bond, the fragmentation is probably inadequate. Using this concept, we extended the fragment-and-cap scheme proposed by [38] by considering resonance via WBOs. The fragment-and-cap scheme generates minimal torsion fragments that include the four torsion atoms and the relevant neighboring atoms that define the local chemical environment [38]. While the scheme captures the local chemical environment, it sometimes does not adequately capture through-bond non-local effects, specifically when extended conjugated systems are involved. We extend the scheme to include the remote substituents that contribute to the electronic overlap across the central torsion bonds by using the WBO as indicators of how well the chemical environment was preserved.

However, WBOs are conformation-dependent [34] which can result in different fragments if the conformations change.Therefore, we use the Electronically Least-Interacting Functional groups (ELF10) WBOs, a method available in the OpenEye toolkit quacpac [qua] that generates WBOs that are relatively insensitive to the initial conformation. But since conformation-dependent variance of WBO distributions are higher for conjugated bonds, we can then use the conformation-dependent distribution of WBOs over conformation to asses whether the chemical environment has been significantly perturbed as described in more detail in section 3.3.

The scheme, illustrated in ***Figure 3*** is as follows:

**Figure 3.**
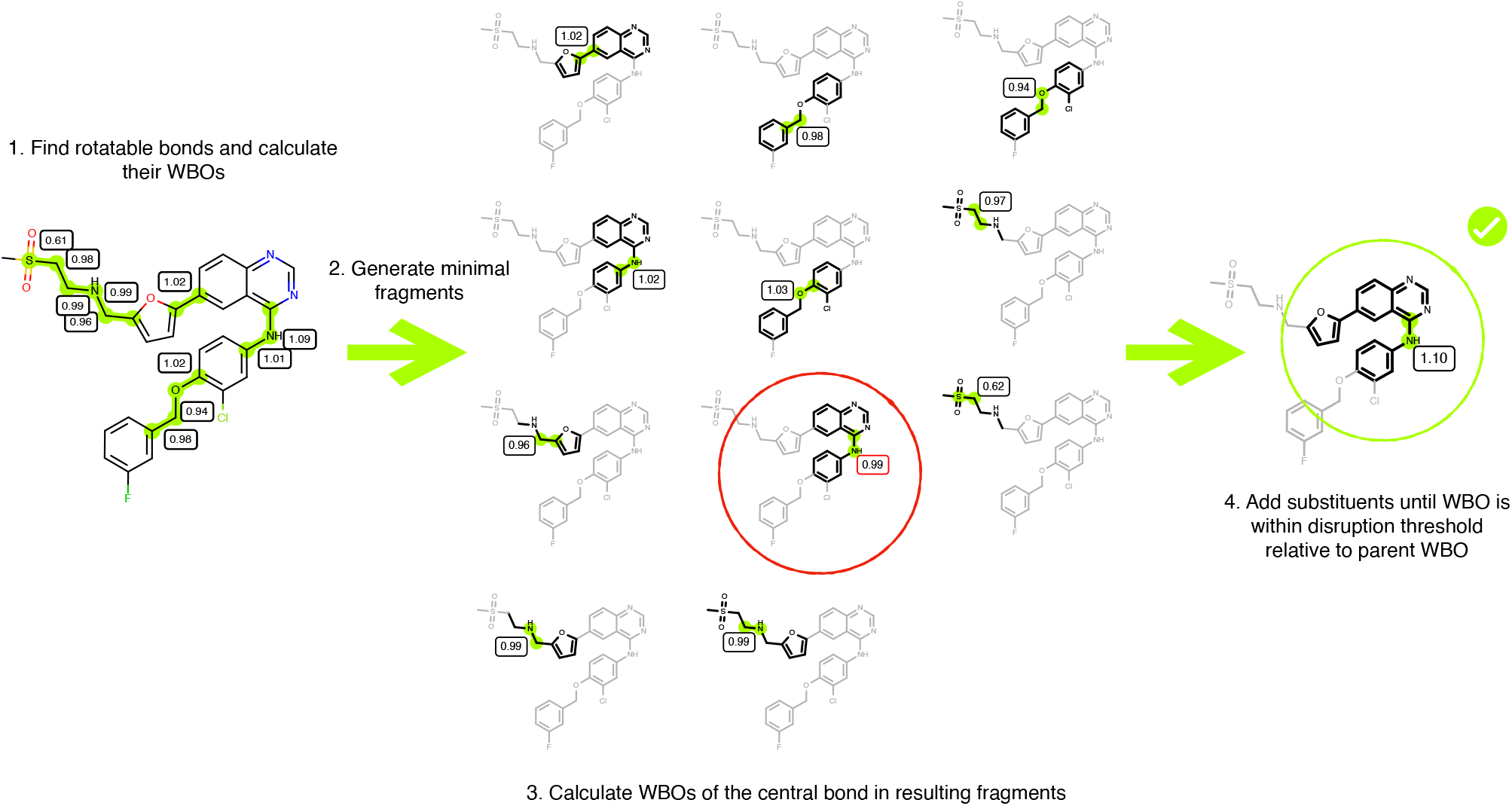
Illustration of fragmentation scheme using WBOs. First, we find the rotatable bonds and calculate their ELF10 WBOs. Then, for each rotatable bond, we generate the minimal fragment about each rotatable bond as described in the text (Section 3.4). We then compute ELF10 WBOs for the central torsion bonds around which the minimal fragments were generated and check if the new ELF10 WBO is within the disruption threshold relative to the parent’s ELF10 WBO. If the absolute difference is greater than the disruption threshold, substituents are added, one at a time, and the ELF10 WBO is recalculated. The red circle illustrates such a fragment. The absolute difference of its WBO relative to the parent (0.99 in the minimal fragment vs 1.09 in the parent molecule) is greater than the disruption threshold of 0.03. The fragment continues to grow until the central bond’s ELF10 WBO is within the disruption threshold of the ELF10 WBO of the bond in the parent molecule. The green circle illustrates the better, larger fragment that has an ELF10 WBO on its central bond closer to the ELF10 WBO on the same bond in the parent molecules.

1. Find acyclic bond. For this step we use the SMARTS pattern [!$(*#*)!D1]-,=;!@[!$(*#*)&!D1].
2. Keep the four atoms in the torsion quartet and all atoms bonded to those atoms. This ensures that all 1-5 atoms are included in the minimal fragment.
3. If any of the atoms are part of a ring or functional group shown in Table 1, include ring and functional groups atoms to avoid ring breaking and fragmenting functional groups that contain more than one heteroatom. The list in Table 1 are the functional groups that were present in the validation and is therefor not exhaustive.
4. Keep all ortho substitutents relative to the torsion quartet atoms.
5. N, O and S are capped with methyl. All other open valence atoms are capped with hydrogen.
6. Calculate ELF10 WBO for fragment.
7. If the fragment’s ELF10 WBO differs by more than a user defined threshold, continue grow out one bond at a time until the fragment’s ELF10 WBO is within the threshold of the parent ELF10 WBO.

### 3.3 Fragmentation schemes can be assessed by their ability to preserve the chemical environment while minimizing fragment size

The fragmentation scheme presented in section 3.2 elaborates upon the method proposed in Rai et. al [38] to remedy deficiencies that will be addressed in more detail in section 3.3.2. However, there are some adjustable hyperparameters. In order to asses various thresholds and different fragmentation schemes in general, we generated a diverse subset of FDA-approved drug molecules that can be a useful validation set. The goal of this set was to find molecules that are challenging to fragment, such that the molecules in this set have bonds that are sensitive to remote substituent changes. In other words, we sought to create a set which is particularly difficult for conventional techniques to fragment well in order to test our method. To find these molecules, we first filtered DrugBank (version 5.1.3 downloaded on 2019-06-06) [51] with the following criteria:

1. FDA-approved small molecules
2. Largest ring size has less than 14 heavy atoms
3. Smallest ring size has at least 3 heavy atoms
4. Molecule has less than 10 rotatable bonds
5. Molecule must have at least one aromatic ring
6. Molecule has only one connected component

This left us with 730 small molecules. Representative molecules are shown in ***Figure 4***. Charged molecules exacerbate remote substituent sensitivity and many molecules are in charged states at physiological pH. To ensure that our dataset is representative of drugs at physiological pH, we used the OpenEye `EnumerateReasonableTautomers` to generate tautomers that are likely significantly populated at pH 7.4 [qua]. This tautomer enumeration extended the set to 1289 small molecules. We then generated all possible fragments of these molecules by using a combinatorial fragmentation scheme. In this scheme, every rotatable bond is fragmented and then all possible connected fragments are generated where the smallest fragment has 4 heavy atoms and the largest fragment is the parent molecule. This scheme generated 300,000 fragments.

**Figure 4.**
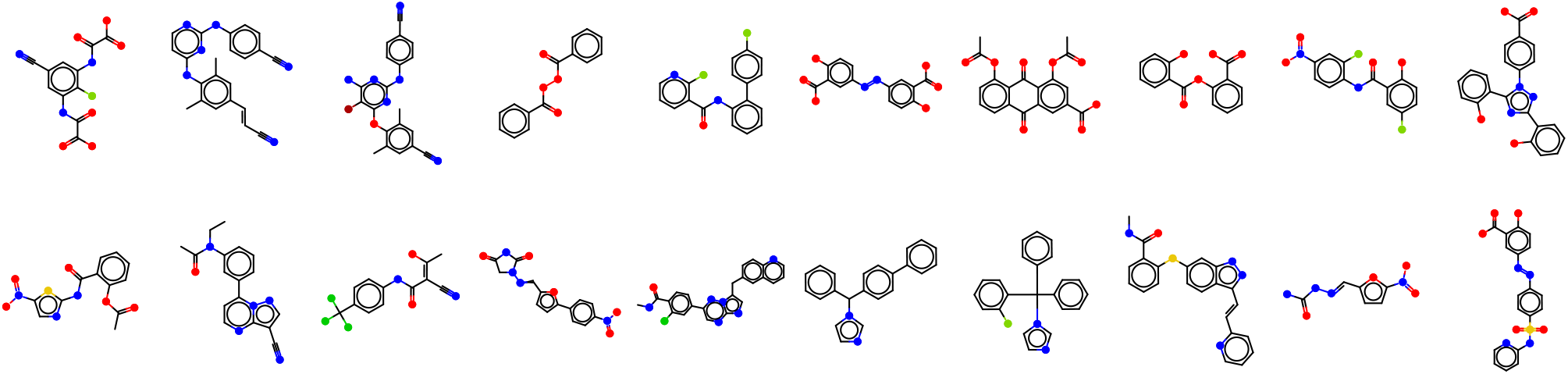
Representative diverse drug-like molecules used to generate the validation set. Twenty representative molecules in the validation set after filtering DrugBank shown here to illustrate the diversity of the set.

To asses how much the chemical environments were perturbed in these fragments, we generated conformation-dependent WBO distributions for every fragment and compared them to the parents’ WBO distributions. Using the conformation-dependent WBO distributions rather than ELF10 WBOs for assessment allowed us to more fully capture the bonds’ chemical environment because changes in variance are also indicative of change in conjugation [42]. For each fragment, OpenEye Omega [20] was used to generate conformers and the AM1 WBO was calculated for every bond in every conformer. This resulted in a distribution of WBOs for all bonds in all fragments. The resulting dataset captures both obvious and subtle through-bond non-local chemical changes.

***Figure 5*** shows an example of the results of exhaustive fragmentation and how this data can be used to benchmark fragmentation schemes. The parent molecule, Dabrafenib (***Figure 5***, C 4) was fragmented at all rotatable bond which resulted in 11 fragments (in this example, the trimethyl was not fragmented). Of these fragments, all connected combinations were generated resulting in 108 connected fragments. Of those 108 fragments, 44 fragments contained the bond between the sulfur in the sulfonamide and phenyl ring highlighted in fragments in ***Figure 5***, C.

**Figure 5.**
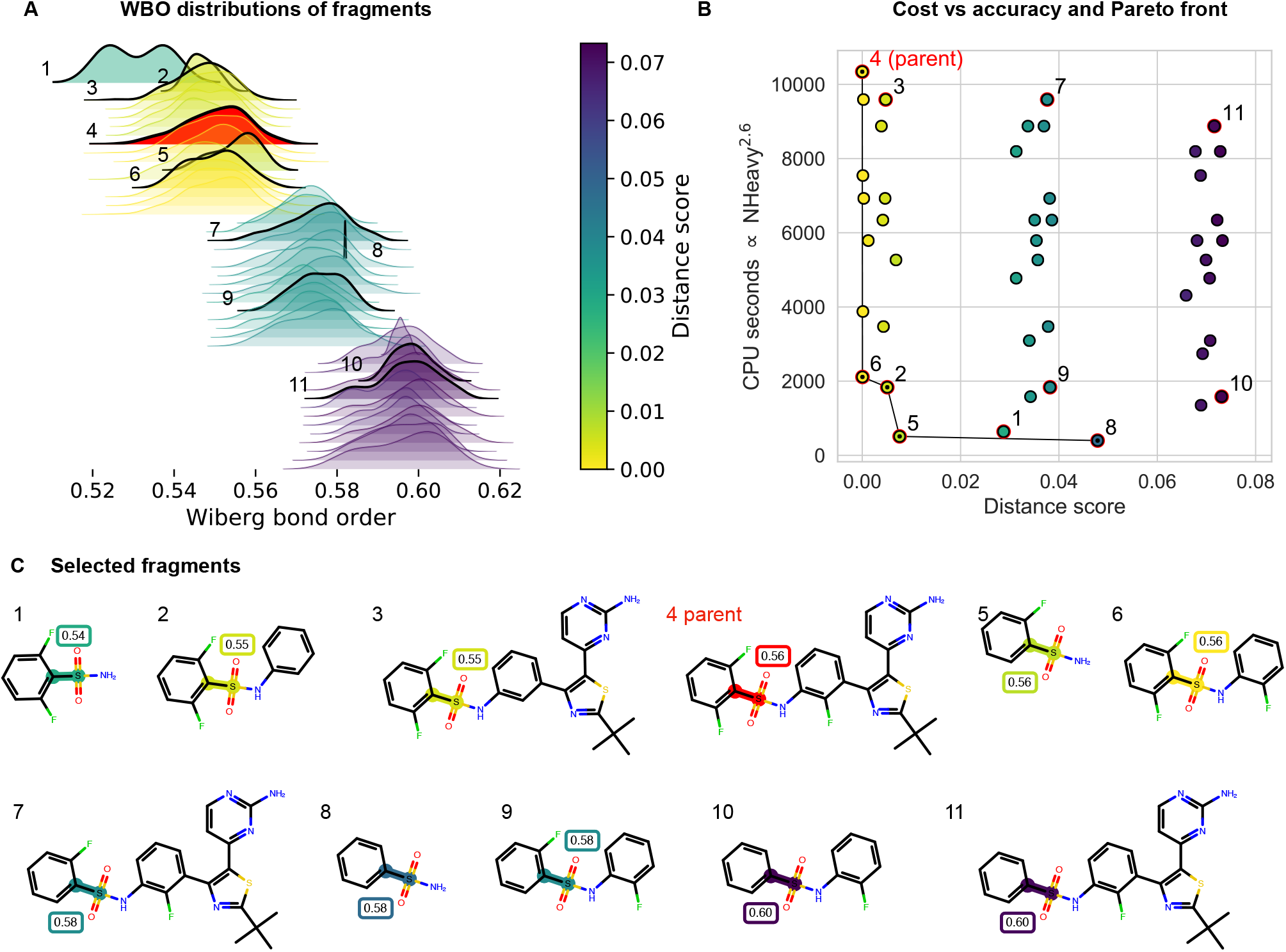
Truncating critical chemical substituents leads to significant changes in the WBO distributions. **[A]** An illustrative example of the shift in the conformational-dependent WBO distributions due to crucial chemical changes such as the loss of fluorine. The distributions are shaded with their corresponding distance score shown in the color bar on the right. The parent molecule’s WBO distribution (numbered 4) is shaded red. Selected distributions are outlined and the corresponding fragments are shown in **C. [B]** Computational cost of fragment (*NHeavy*^2.6^) vs distance score (Maximum Mean Discrepancy, Equation 3.3.1) of the fragment indicates that it is possible to reduce the cost of torsion scans without destroying the torsion profile. The black line is the Pareto front, or the most computational inexpensive fragment with the best score at that size. The selected fragment should be on the Pareto front at the lower left corner. **[C]** Selected fragments. Bonds are highlighted with their distance score. The ELF10 WBO is shown in the boxes above the highlighted bonds.

When the WBOs were calculated for all Omega generated conformers for each of the 44 fragments, the resulting WBO distributions clustered into 4 bins as illustrated with the color and location of the distributions in ***Figure 5***, A. The shifts of the distributions corresponded to specific remote substituent changes—in this case, the loss of fluorine of the ring bonded to the sulfur and the phenyl ring bonded to the nitrogen in the sulfonamide. Excluding the flourine on the other ring (bonded to the nitrogen) does not influence this metric as seen by the significant overlap between the WBO distribution of the parent and fragment 3 in ***Figure 5*** A. Here, these two changes cause the distributions to shift in opposite directions. While the loss of a fluorine on the phenyl bonded to the sulfur shifts the distribution to the right, the loss of the ring bonded to the nitrogen shifts the distributions to the left illustrating that the changes can counteract each others. Fragments 2, 3, 4, and 6 (***Figure 5***C) all contain two fluorine and the phenyl bonded to the nitrogen and fall in the same cluster as the parent molecule, regardless if the rest of the molecule is included in the fragment. Fragments 7 and 9 only have one fluorine on the phenyl ring and both of their distributions are shifted to the right relative to the parent WBO distribution. Fragments 10 and 11 have no fluorine on the ring and are shifted to the right even further. Since removing the ring bonded to the nitrogen shifts the WBO distribution in the opposite direction, fragment 1, while having two fluorine, is shifted to the left of the parent distribution, fragment 5 WBO distribution overlaps with the parent WBO distribution even if it only has one fluorine, and fragment 8 is only shifted slightly to the right of the parent WBO distribution with no fluorine.

#### 3.3.1 WBO distributions can measure how well fragments preserve the chemical environment around the torsion

Each fragment needs to be assigned a score of how well is preserves its parent chemical environment. To score each fragment, we compare the conformer dependent WBO distribution for a bond in a fragment against the WBO conformer-dependent distribution of the same bond in the parent molecule. To compare these distributions, we compute the maximum mean discrepancy (MMD) [14] fragment distribution to the parent as follows:

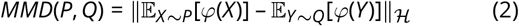

where the feature map *φ* : 𝒳 → ℋ we use is squared *φ*(*x*) = (*x, x*^2^) and the MMD becomes:

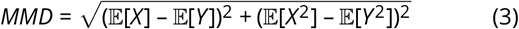

where 𝔼 [*X*] is the expected value of the parent WBO and 𝔼 [*Y*] is the expected value of fragment WBO. Including the squared mean incorporates the second moment of the distribution and helps distinguish distributions both with different means and variances. It is important to incorporate changes in variance given how the variance of the WBO distributions change for different chemical environments [42]. In general, changes in WBO distribution variance corresponds to changes in relative torsion barrier heights [42].

In ***Figure 5***, the MMD, which we refer to distance score, is shown with the color map. The distributions in ***Figure 5***, A are shaded with the distance score. The scores clearly differentiates the shifted distributions.

#### 3.3.2 Good fragmentation schemes minimize both chemical environment disruption and fragment size

The goal of our fragmentation scheme is to find fragments that have a WBO distribution of the bond of interest closest the the parent while minimizing the computational cost of the fragment. We estimate the computational cost of a fragment as *NHeavy*^2.6^ as shown in ***Figure 1*** A. The distance score calculated with MMD indicates how far the fragment’s WBO distribution is or how much the chemical environment changed from its parent. When we plot the fragment size against this score, the points that fall on the Pareto front [30] are the ones where the distance score is the best for for a given fragment size or vice versa.

***Figure 5***, B shows an illustrative example of this. The Pareto front is shown with a black line and the fragment data points on the front are indicated with a black circles. The numbers on the annotated data points correspond the the numbered fragments in ***Figure 5***, C. Fragment 6 is the smallest fragment with the shortest distance to the parent molecule. It has the important chemical moieties, such as all three fluorine and the ring bonded to the nitrogen. While fragments 2 and 5 are also on the Pareto front, the missing ring and fluorine increase the distance score, however, it is not clear if this difference is significant.

Fragment 3, which is missing the fluorine on the ring bonded to the nitrogen, is shifted in the distance score relative to the parent by the same amount as fragment 2 is from 6, even if it has all other parts of the molecule. Both the chemical change and shift in distance scores are small, but they are consistent for both larger and smaller fragments. adding credence to the fact that the small difference in the distance score does pick up on this chemical change. Fragments 7 and 11 illustrate that having larger fragments will not improve the distance score if the important remote substituents are not present in the fragment. Fragment 9, while significantly smaller than fragment 7, has the same distance score because they both are missing the important fluorine. Fragments 10 and 11 show the same trend for the fragments missing both fluorine. While fragments 1, 5, and 8 are all small, the loss of the ring results in larger distance scores.

The goal of any fragmentation scheme is to find fragments on the Pareto front that minimize both the changes in the chemical environment of the bond and fragment size. In other words, they should be on the lower left corner of the plot.

To assess the quality our fragmentation scheme, we wanted to find the molecules that are challenging to fragment. To do that, we defined a bond sensitivity score as follows. For each bond, find all fragments from the exhaustive fragmentation experiment that include the bond, calculate the distance score to the parent, and find the fragment with the maximum distance. This maximum distance score is the sensitivity score of the bond. This is a good indication of a bond’s sensitivity to removal of remote substituents because the more its WBO distribution shifts relative to the parent when fragmented, the more the electronic population overlap around that bond changes with remote chemical changes. We then chose the top 100 molecules that had bonds where fragments that included all 1-5 atoms around the central bond had the highest sensitivity scores Selected molecules with bonds highlighted according to their sensitivity are shown in ***Figure 6***. The rest of the molecules are shown in SI ***Figure 5***. This set included many molecules in charged states.

**Figure 6.**
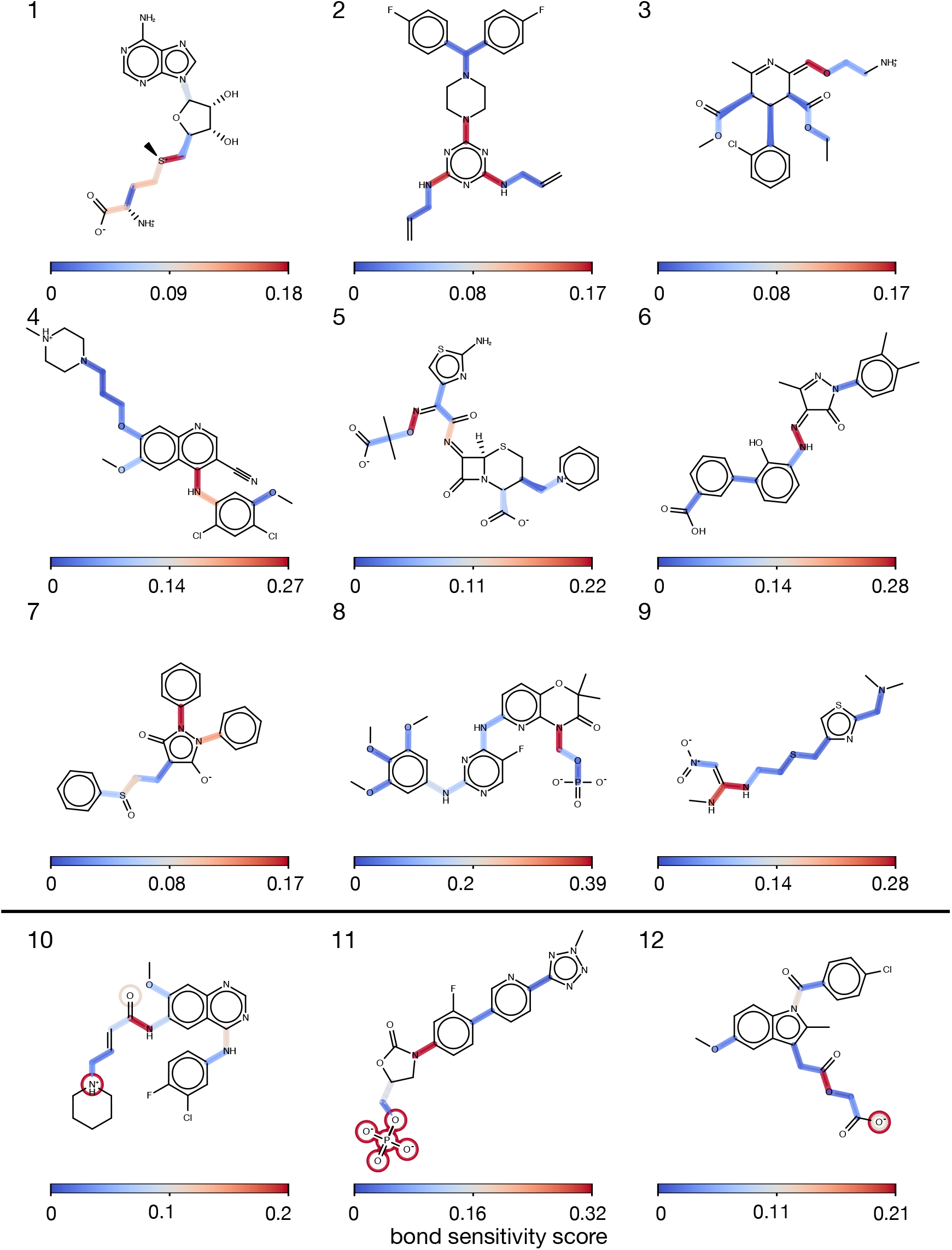
Some bonds are more sensitive to removal of remote substituents and are therefore challenging to fragment. Selected molecules of the validation set are shown to illustrate differential bond sensitivity to remote substituent changes. Bonds are highlighted by how sensitive they are to remote substituent changes as quantified by the bond sensitivity score. The bond sensitivity score is defined as the maximum distance score of that bond in different fragments as defined in the text. (see ***Figure 5*** A and B). Fragments used to get the maximum distances include all 1-5 atoms around the highlighted bonds. Molecules 10-12 also show the atoms the bonds are sensitive to. The atoms are circled with the same color as the bond that is sensitive to it. The rest of the molecules used in the validation set are shown in SI ***Figure 5***

Not all bonds are equally sensitive to such changes. This is shown by the magnitude by which the sensitivity score varies for bonds within the same molecule as shown in ***Figure 6***. The general trend observed is that conjugated bonds, and bonds including atoms with lone pairs such as N, O, and S, are more sensitive to peripheral cuts. Molecules 10-12 (***Figure 6***) also show which chemical moiety the bond is sensitive to, indicated by circles around the atoms which are colored with the corresponding bond’s sensitivity score. In molecule 10, the WBO distribution of the amide bond shifts significantly if the positively charged nitrogen is remove regardless if the rest of the molecule is intact (data not shown). In molecule 11, the removal of the phosphate group shifts the distribution of the red bond. In molecule 12, both the amide and ester bond are sensitive to the same negatively charged oxygen indicated by two circles around the oxygen.

We aim to identify parameters for our fragmentation scheme that maximize the number of fragments that end up in that lower left corner of the Pareto front (illustrated in ***Figure 5***, B). To do that, we generated fragments for the highly sensitive bonds in the 100 molecules shown in SI ***Figure 5*** using different disruption thresholds. For every fragment, we found the distance score of their fragments’ WBO distribution and their computational cost. We then plotted all fragments from the validation set for different thresholds (***Figure 7***). When the threshold is low, the fragmentation scheme will generate fragments which have very good distance scores, but many of them will be too big for computationally efficient QC torsion scan. On the other hand, when the disruption threshold is too low, the scheme generates fragments that are small but the distance scores are too big. For the molecules we tested, a threshold of 0.03 leads to the most fragments in the lower left quadrant (defined as cost < 4000 seconds per gradient evaluation and score < 0.05) as shown in Table 2.

**Figure 7.**
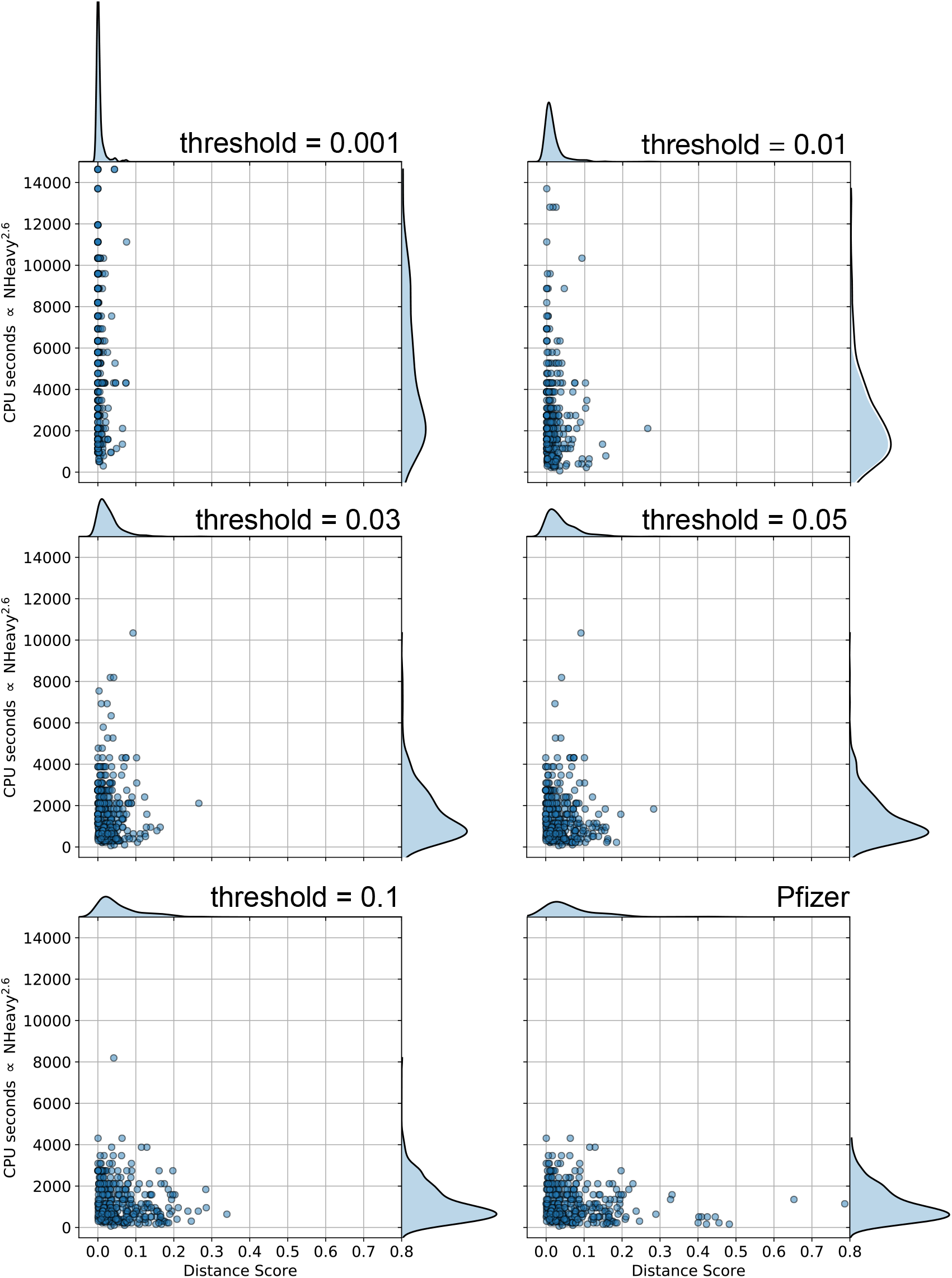
Different WBO disruption thresholds result in different accuracy vs cost trade-offs. Computational cost vs distance score of the fragments for the bonds in the benchmark set shown in ***Figure 5***. Computational cost is proportional to *NHeavy*^2^.6 as estimated in ***Figure 1***, A. The threshold is the maximum amount of change allowed in the ELF10 WBO relative to the parent molecules’s ELF10 WBO during fragmentation.

**Table 2.** Number of fragments in the lower left quadrant in ***Figure 7*** defined as a distance score less than 0.1 and computational cost less than 4000 seconds.

| WBO disruption threshold | Number of fragments in optimal fragment quadrant |
| --- | --- |
| 0.001 | 182 |
| 0.005 | 232 |
| 0.01 | 268 |
| 0.03 | 284 |
| 0.05 | 243 |
| 0.07 | 208 |
| 0.1 | 209 |
| Pfizer scheme [38] | 197 |

This threshold is similar to what we found when we looked at the distribution of standard deviations for WBO distributions with respect to conformations [42]. Most of them fall under 0.03. Both of these data points lead us to recommend a WBO disruption threshold of 0.03 for our fragmentation scheme. While the current scheme does not provide a perfect solution, plots in ***Figure 7*** shows fewer fragments outside of the lower left region for thresholds 0.01, 0.03 and 0.05. This scheme performs better than other schemes such as the scheme Pfizer used in [38] (***Figure 7***, lower right and Table 2).

#### 3.3.3 Benchmark results reveal chemical groups that induce long range effects

In the benchmark experiment (***Figure 7***), Omega was used to generate the conformers for which the WBOs were calculated. Omega aims to generate low energy conformers [20] and in some cases, fragments only have one or two low energy conformers so it is not clear how accurate the distances measured are. In addition, only comparing low energy conformers do not fully capture torsion energy barriers which we also want to ensure remain accurate relative to their parent’s torsion energy scan.

To mitigate these issues when validating our scheme, we generated higher energy conformers by driving the torsions and added those WBOs to the distributions. Furthermore, since we know that WBOs from structures in a QC torsion scan are anti correlated with the QC torsion energy scan [42], adding WBOs from higher energy conformers to the distributions provided a better validation of our method than examining only the distance between WBOs from low energy omega-generated conformers. The differences in distances of these distributions from fragments generated with our scheme and the scheme in Rai et. al [38] is shown in ***Figure 8***.

**Figure 8.**
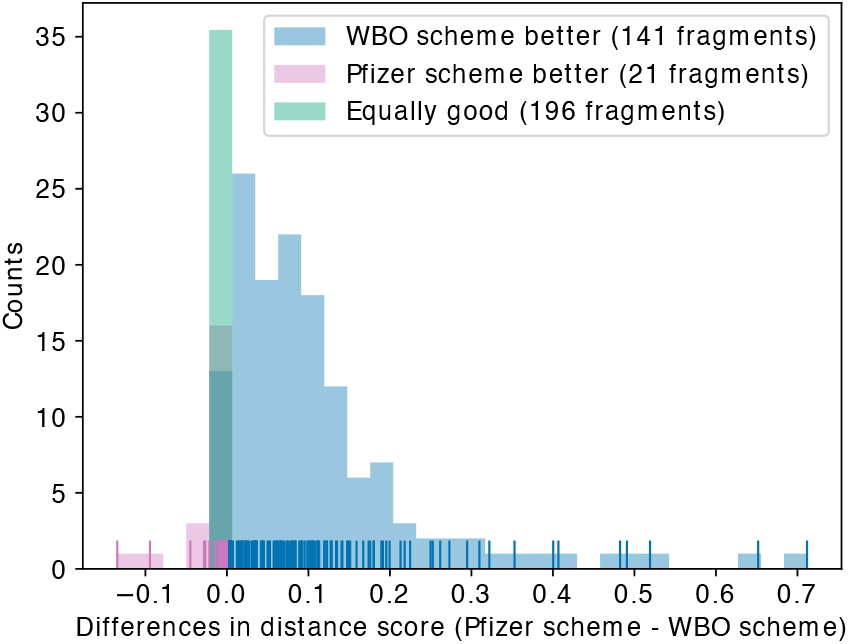
Using the WBO as an indicator of chemical environment disruption improves performance of the fragmentation algorithm. Distribution of differences in distance scores for fragments in the validation set (SI ***Figure 5***) generated using Pfizer’s rules and our scheme using 0.03 as the disruption threshold. For many bonds, both approaches yield equally performing fragments (shown in green). In some cases, Pfizer’s rules [38] perform better than our scheme (shown in red); however, the differences are usually very small. In most cases, using the WBO as an indicator improves the distance score (shown in blue).

For many molecules, using a common sense rule based approach, such as the one used in Rai et. al [38] to fragmenting molecules, will yield the same fragments our scheme generated. The equally good fragments are shown in green in ***Figure 8***. Sometimes a simple approach can even perform slightly better than using the WBO as an indicator (***Figure 8***, red). However, in some cases, especially if certain chemical groups are involved, using the WBO as an indicator significantly improves the electron population overlap about the bonds and brings them closer to their parent’s chemical environment (***Figure 8***, blue). It is important to note that when the fragment generated from both scheme are the same (green in ***Figure 8***), they are not necessarily the optimal fragment and both schemes can perform equally poorly (SI ***Figure 6***, A, B, and C). However, in most cases both fragments do perform well (SI ***Figure 6***, D, E, and F).

Upon closer inspection of the validation set, we found eight chemical groups that induce long range effects to sensitive bonds. These chemical groups with representative examples are shown in ***Figure 9***. The groups are ordered by how strongly they induce long range effect, in decreasing order. The most dramatic change happens when a phosphate group is removed (***Figure 9***, A). The variance of the WBO distribution increases which conveys an increase in relative energies of conformers in the QC torsion scans. In other molecules where phosphates are removed, the variance can decrease even if the phosphate group is ten bonds away (***Figure 10***, F and SI ***Figure 7*** E).

**Figure 9.**
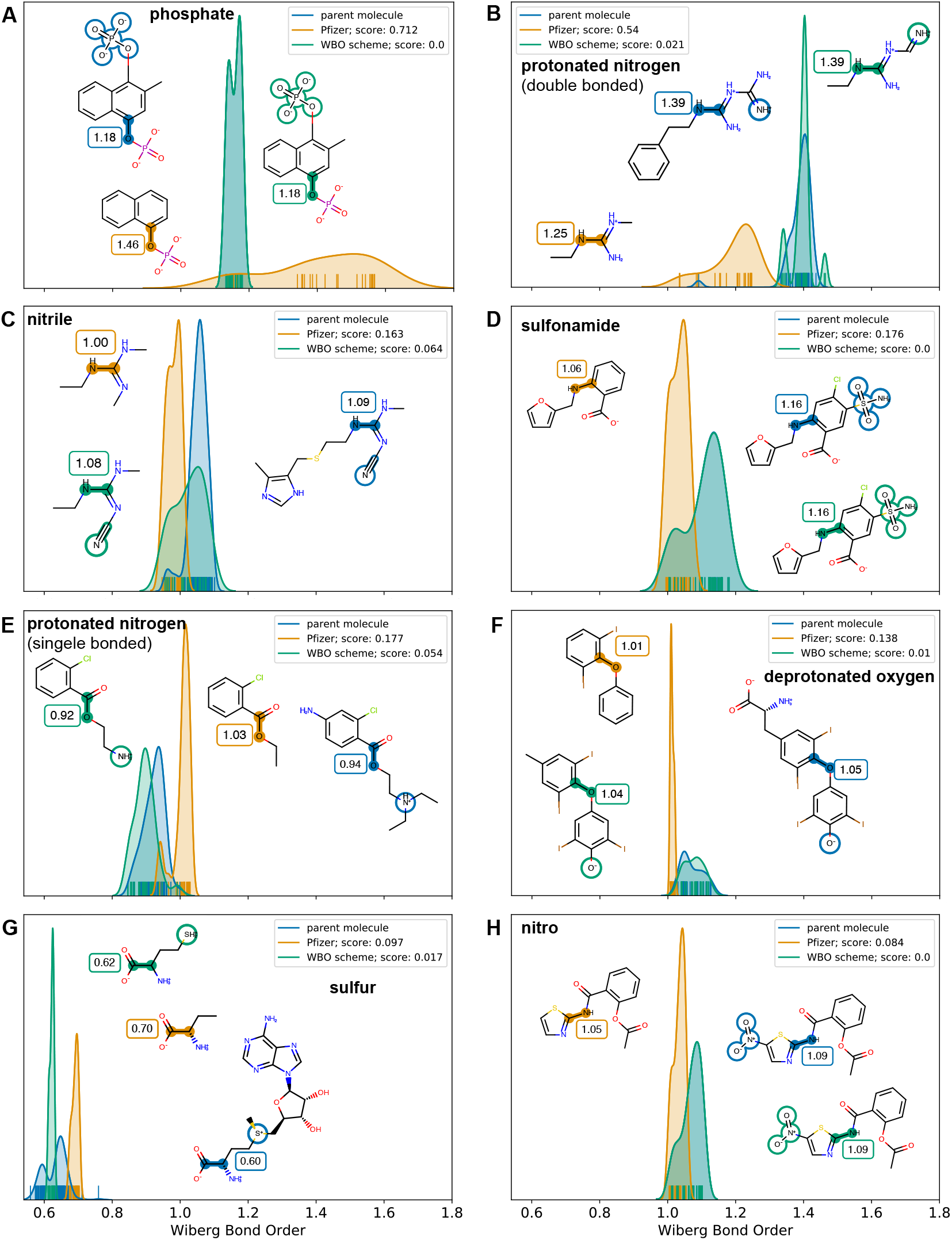
Some chemical groups induce non local effects that are captured in fragments when using the WBO as an indicator of chemical environments. Wiberg bond order distributions for parent molecules (shown in blue) and fragments generated with Pfizer rules (shown in orange) and our scheme (shown in green). The distribution is over all Omega generated conformers and high energy conformers from torsion drives. The rug plot at the bottom of the distributions are tick marks for each observation. This figure shows eight chemical groups where the WBO distributions of the highlighted bonds change when those groups are removed. The ELF10 WBO is shown in the boxes next to the highlighted bonds and the chemical groups are circled in the parent and our fragment. Better overlap with the blue, parent distribution, indicates a fragment where the chemical environment around the highlighted bond is closer to the parent environment. In the examples shown here, our scheme generates fragments where the WBO distribution of the bond of interest has better overlap with the parent WBO distribution. Changes are consistent across the validation set. Since the parent molecule in **[G]** has several rotatable bonds, there are many tick marks in its rug plot.

**Figure 10.**
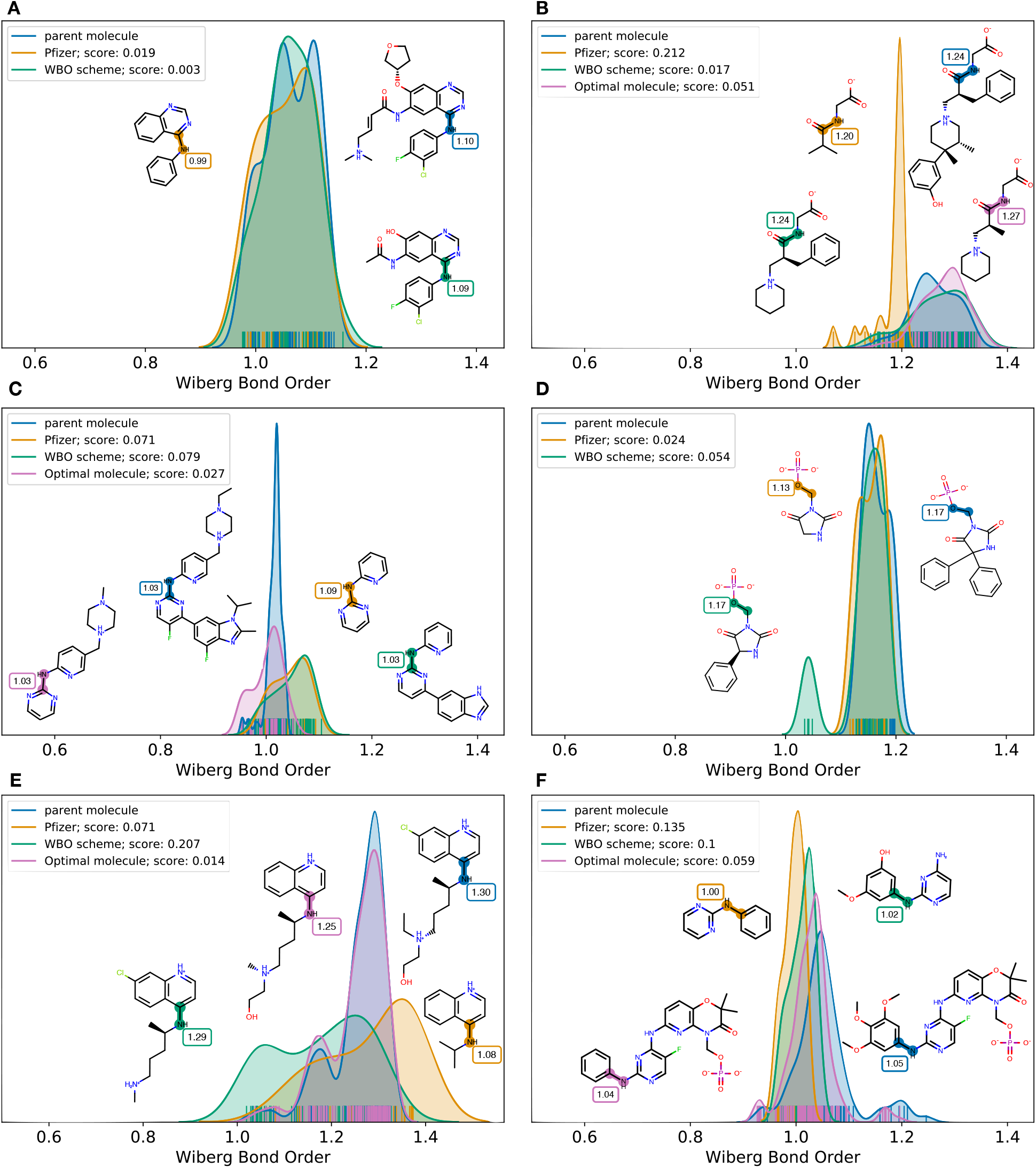
Using the WBO as an indicator when fragmenting can still fail to find the optimal fragment. Our scheme can fail in several ways. **[A]** A smaller fragment (shown in orange) is just as good as a larger fragment (shown in green) even if the ELF10 WBO estimate of the bond in the smaller fragment relative to its parent (shown in blue) is more than the disruption threshold. **[B]** While our scheme finds a fragment with good overlap of the WBO distributions (shown in green), it is not the smallest fragment possible with good distributions overlap (smallest fragment with good overlap is shown in purple). **[C]** The fragment we find is bigger than what the simple scheme finds (shown in orange) but without improving the WBO overlap (green). The optimal fragment that neither scheme generates is shown in purple. **[D]** Our scheme finds a larger fragment that has worse WBO distribution overlap. **[E]** and **[F]**. Sometimes, almost the entire molecule is needed to achieve good WBO distribution overlap between the fragment and the parent. This is not a failure mode but inherent to the challenge of fragmenting molecules for QC calculations.

In ***Figure 9***, B, removing a protonated nitrogen that is double bonded causes the WBO distribution to shift and the variance to increase. Long range effects are seen in other molecules with similar chemical patterns up to eight bonds away (SI ***Figure 8***, E). Removing a nitrile group (***Figure 9***, C) and sulfonamide group (***Figure 9***, D) have similar effects on the WBO distributions which is also consistent with other molecules that contain these groups up to three bonds away (SI FIGsi:si-nitrile and ***Figure 10***). A protonated nitrogen and deprotonated oxygen (***Figure 9*** E and F) can effects bonds between 3-6 bonds away (SI ***Figure 8*** and ***Figure 11***). While the changes in distributions for removing a nitro group and sulfur (***Figure 9***, G and H) are not as big as other chemical groups, they are mostly consistent across other molecules in the validation set (SI ***Figure 12*** and ***Figure 13***).

While our scheme captures long range effects that a simple rule based approach does not, it is not an optimal solution and will sometimes fail to find the most optimal fragment. By optimal we mean the smallest fragment that retains the torsion potential of the bond in the parent molecule. Our scheme can fail in multiple ways as illustrated in ***Figure 10*** and listed below.

1. The selected fragment has good WBO distributions overlap but is not the smallest fragment. This is illustrated in both ***Figure 10*** A and B. In A, the fragment that Pfizer scheme find is smaller and has a WBO distribution that is close to the parent’s WBO distribution (MMD 0.019). In this case, the ELF10 WBO estimate of the bond in the fragment is 0.11 lower than the ELF10 WBO estimate in the parent molecule. In B, our fragment has better WBO distribution overlap with the parent WBO distribution vs using Pfizer’s scheme (0.017 vs 0.212), but it is not the smallest fragment. According to the fragment highlighted in purple, the benzene ring is not required to achieve good overlap of the WBO distributions (0.051).
2. Sometimes, the fragments we find are bigger than Pfizer’s scheme fragment and the remote substituents do not improve the WBO distribution overlap (MMD 0.079 vs 0.071) (***Figure 10*** C). The better fragment is shown in purple. It is both smaller and has better overlap (MMD 0.027) than the orange and green fragment.
3. Sometimes, the fragment we find is both larger and has worse overlap (***Figure 10***, D; 0.054 vs 0.024) than what the Pfizer’s scheme generates.
4. While it is usually possible to find a fragment that is significantly smaller than the parent and retains remote substituent effects, the effects are sometimes more than 3-6 bonds away and a large fragment is needed to accurately represent the chemical environment of the parent molecule. Two such examples are shown in ***Figure 10*** E and F. In E, not only is the protonated nitrogen needed (shown in green), but the alcohol group is also needed to achieve good WBO distribution overlap (shown in purple). In F, the phosphate group nine bonds away from the bond of interest is needed to capture the density of of the mode at 1.2 (shown in blue and purple).

As we have shown, using the WBO as an indicator for fragmentation is a good general approach, albeit not always optimal. To always find the optimal fragment, all possible fragments would need to be generated and their WBOs calculated. As discussed in section 2.1, the search space for all possible fragments can become very large which would be too costly to enumerate. Therefore, we chose to use heuristics discussed in the detailed methods (5.2.3) to grow out the fragments when the WBO on the central bond indicates that the chemical environment has been disrupted.

#### 3.3.4 Validating the fragmentation scheme

To validate our fragmentation scheme, we ran several series of torsion scans on parent molecules and fragments - both fragments generated using only the rules used in Rai et. Al [38] and fragments using our WBO based scheme. For all the molecules we tested, we found that the torsion scans of the fragments using the different schemes differed significantly as shown in ***Figure 11*** To avoid high computational cost and through-space intramolecular interactions, we chose parent molecules that were less than 21 heavy atoms. Therefore, some of the fragments found by our WBO scheme were the parent molecules (***Figure 11*** D and F). For the cases where the WBO scheme generated fragments that were smaller than the parent molecules, the QC torsion scan of those fragments were closer to the QC torsion scans in the parent molecule than for the fragments generated with simple rules. The only exception is ***Figure 11*** E where the QC scan of the fragment without the nitrile has a smaller RMSE relative to the parent QC scan as the WBO generated fragment (RMSE 6.38 vs 6.97). This discrepancy is a result of the asymmetry in the parent’s molecule QC torsion scan (blue scan). The asymmetry arises from favorable through-space interactions between the nitrile and hydrogens on the methyl on the imidazole at dihedral angles 15*°* - 115*°*. Such long-range through-space favorable interactions are effects that should be handled by non-bonded force field terms so they should be avoided in the data used to fit torsion parameters to. Therefore, we view ***Figure 11*** E as a good result.

**Figure 11.**
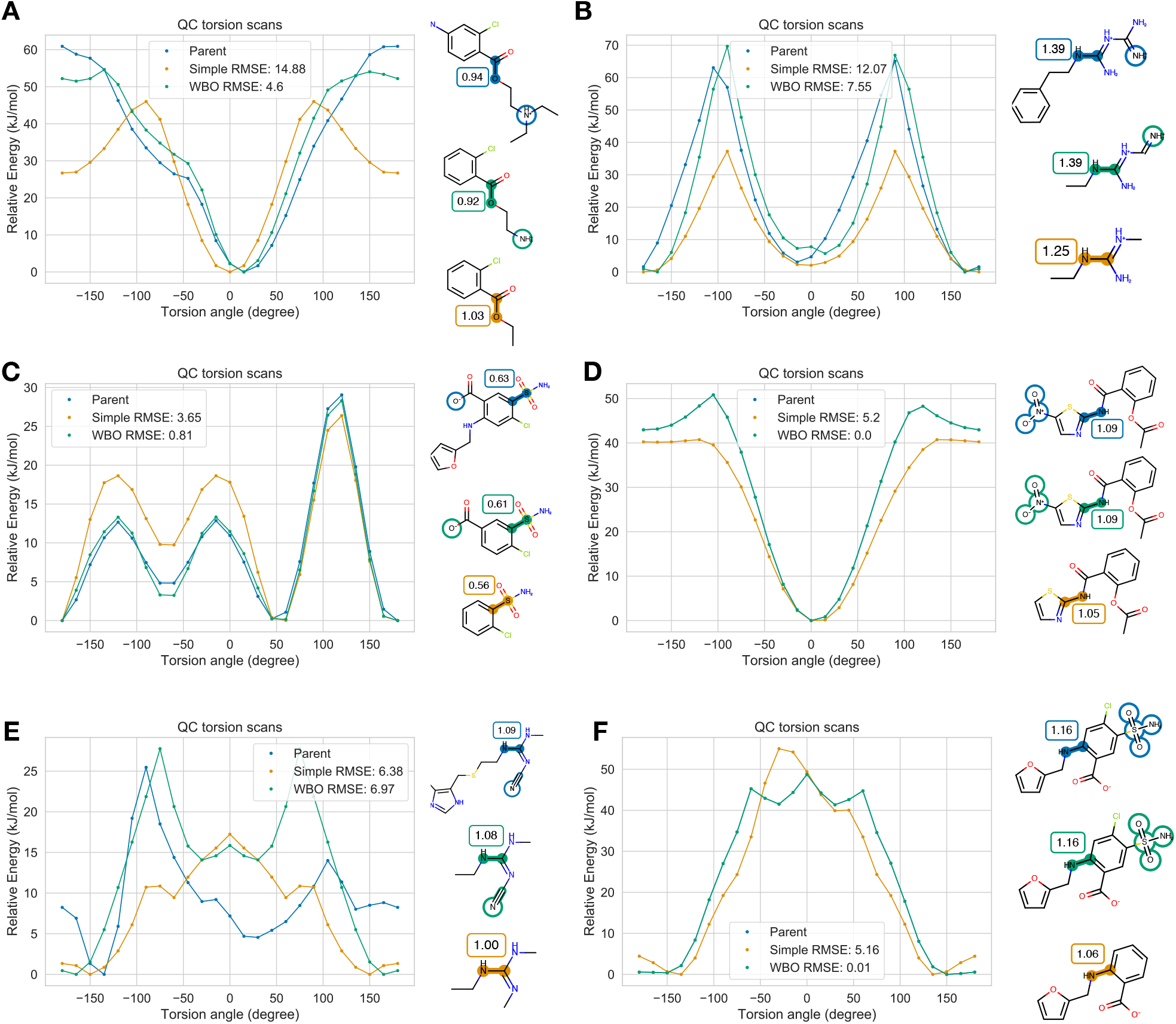
Using WBO as a criteria for fragmentation improves the chemical equivalency of the fragment to the substructure in the parent molecule. Representative QC torsion profiles of drug molecules (parent molecules shown in blue), fragments generated from rule based scheme (shown in orange) and fragments generated with the WBO scheme (green). **[A]** Torsion profile about the highlighted bond in Chloroprocaine for the parent molecule (blue), WBO scheme (green) and simple rules (orange). **[B]** Same as **A** for Phenformin. **[C]** Same as **A** for Furosemide. **[D]** Same as **A** for Nitazoxanide. Here, the parent and fragment generated with the WBO scheme are the same. **[E]** Same as **A** for Cimetidine. **[F]** Same as **C** for Furosemide.

## 4 Conclusion

We have shown that the AM1 ELF10 WBO estimate is a simple, yet informative quantity about the extent of binding between two connecting atoms, thus descriptive of a bond’s chemical environment, its level of conjugation, and resistance to rotation. We can use the change in WBO of a bond to quantify the amount of disruption of its chemical environment due to remote chemical substituent changes, specifically for bonds that are sensitive to peripheral chemical changes such as bonds in or adjacent to conjugated systems, or bonds involving atoms that have lone pairs.

We used this concept to extend a rule-based fragmentation scheme to improve the resulting fragments. By using the WBO of the central bond as an indicator of its chemical environment disruption, our extension adds remote substituents until the change in WBO is lower than a user defined threshold. This simple addition improves the quality of the resulting fragments, specifically for bonds in conjugated systems and molecules with remote substituents that have long-range effects. We generated a validation set using exhaustive fragmentation to benchmark fragmentation schemes and found that a threshold of 0.03 will find the most fragments that minimize both fragment size and distance to the parent’s conformation distribution about the bond. We also found eight chemical groups that have long-range effects on sensitive bonds and their inclusion is necessary to recapitulate a parent’s chemical environment even if they are 3-6 bonds away from the central bond.

Overall, we believe the approach presented here provides a good general approach to fragment molecules for QC data generation to assist in parametrization of general force fields. On our test set, this approach seems superior to that of straightforward alternatives. The method as described in this manuscript is available as an open source Python library https://github.com/openforcefield/fragmenter. We plan to apply this approach broadly within the OpenFF Initiative. The method has been incorporate in the open force field toolkit https://github.com/openforcefield/openforcefield. In the future, this approach can be made more efficient by using machine learning methods to learn the chemical patterns that should not be fragmented using the dataset shared here.

## 5 Detailed methods

### 5.1 Exhaustive fragmentation dataset

The exhaustive fragmentation dataset was generated by filtering DrugBank version (version 5.1.3 downloaded on 2019-06-06) [51] with criteria described in section 4 and repeated here for clarity:

1. FDA approved
2. Ring sized between 3 and 14 heavy atoms
3. Rotatable bonds between 4 and 10
4. At least one aromatic ring
5. Only a single connected component

This left us with 730 molecules. To expand protonation and tautomeric states, we used `OEGetReasonableTautomers` from QUACPAC (OpenEye ToolKit version 2019.Apr.2) in the `states.py` module in `fragmenter` (version `v0.0.2+175.g6fbbf32` for this original set, but versions v0.0.3, v0.0.4 and 0.0.6 will generate the same results with the same options). We set `pKaNorm` to True so that the ionization state of each tautomer is assigned to a predominant state at pH 7.4. This generated 1289 molecules. The smi file with all 1289 molecules is located in the manuscript’s data repo https://github.com/choderalab/fragmenter_data/blob/master/combinatorial_fragmentation/enumerate_states/validation_set.smi

We then used the `CombinatorialFragmenter` from `fragmenter` version v0.0.2+179.g0e7e9e3 (versions v0.0.3 and v0.0.4 will generate the same fragments with the same options) to generate all possible fragments for each molecules. We set the option `functional_groups` to False so that all functional groups besides rings will also get fragmented so we can use the data to explore which functional groups should not be fragmented. We used the default settings for all other options (`min_rotor` is 1 and `min_heavy_atoms` is 4 so that the smallest fragments have at least one torsion. `max_rotors` is the number of rotors in the parent molecules so that the largest fragment generated is one less rotor than the parent molecule). This generated 300,000 fragments.

We then used Omega (OpenEye version 2019.Apr.2) to generate conformers for each fragment and calculated each conformer’s WBOs as described above. All scripts used to generate this dataset are in https://github.com/choderalab/fragmenter_data/tree/master/combinatorial_fragmentation. The dataset is available on Zenodo [41].

The benchmark set used to evaluate disruption thresholds and compare to other schemes were chosen as described in Section 3.3.2. `fragmenter` version 0.0.6 was used to generate fragments for the different schemes and disruption thresholds.

### 5.2 Fragmenting molecules

The `fragmenter` package provides several fragmentation schemes with various options. Below we discuss different modes of fragmentation and their options.

#### 5.2.1 Exhaustive fragmentation generates all possible fragments of a parent molecule

This functionality is provided by the `CombinatorialFragmenter` class in the `fragment.py` module. To use this class, the user needs to provide an openeye molecule. `fragmenter` provides a list of functional groups SMARTS in a yaml file located in `fragmenter/data/fgroup_smarts_combs.yml` that it will not fragment by default. This list is given in table 3. The list is different than the default list used on the `WBOFragmenter` because here the carbon bonded to the functional groups are also tagged. To allow all functional groups to be fragmented, the user can set the parameter `functional_groups = False`. This option will fragment all bonds besides bond in rings. The user can also provide their own dictionary of functional group SMARTS patterns that they wish to avoid fragmenting.

**Table 3.**
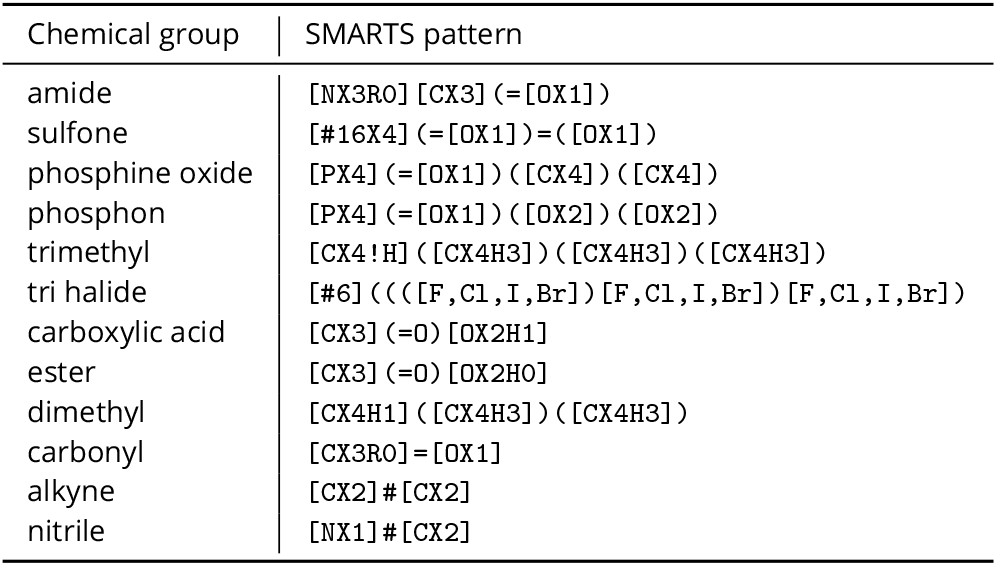
Default functional groups **CombinatorialFragmenter** preserves whole during fragmentation. This list is not comprehensive and is different than the list used for the `WBOFragmenter`.

The user can also set the minimum and maximum number of rotatable bonds, and minimum heavy atoms in a fragment.

#### 5.2.2 Generate minimal fragments

The `PfizerFragmenter` implements the scheme developed at Pfizer and described in Rai et. al [38]. It uses the same list of functional groups as the `WBOFragmenter` uses. The user can also provide their own SMARTS patterns of functional groups not to fragment.

#### 5.2.3 Using the WBO as a surrogate for changes in chemical environment

The `WBOFragmenter` implements the FBO scheme described in this paper. The functional groups that are not fragmented are given in table 1. Users can add more SMARTS patterns if needed.

When the WBO on a central bond in a minimal fragment has been disrupted more than the threshold, remote atoms need to be added onto the fragment. However, there are multiple ways to grow out a fragment and enumerating all possible ways to find the best one can become too computationally expensive. Therefore, we need to use heuristics to decide where to add the next atoms. The two heuristics available in `fragmenter` are:

1. **Shortest path length** Here, the bond with the shortest path to the central bond is added next. The rationale for this heuristic is that atoms closer to the central bond will have a greater influence to the bond’s chemical environment. If more than one connected bond has the shortest path to the central bond, the bond with the higher WBO is added next.
2. **Greatest WBO** Here, the bonds connected to the fragment that have the greatest WBO is added next. The rationale for this heuristic is that bonds with higher WBO are more likely to be involved in extended conjugation that can influence the central bond.

Both of these heuristics will sometimes give different results (SI ***Figure 14***. We found that for the benchmark set we tested, the shortest path heuristic preformed slightly better, or found more optimal fragments when compared to using the greatest WBO heuristic (284 vs 262 in the optimal quadrant). SI***Figure 14*** compares the scores and sizes of fragments generated with different growth paths. The shortest path length heuristic generates smaller fragments (SI ***Figure 14*** A)

### 5.3 Fragmenter validation dataset

The torsion scans shown in ***Figure 11*** were run on QCArchive [39]. They were computed with TorsionDrive [37] and geomeTRIC [47], as standalone geometry optimizer interfaced with the QCEngine [@url:https://github.com/MolSSI/QCEngine] project. The gradients were computed at the B3LYP-D3(BJ)/DZVP level of theory [2, 10, 16] with Psi4 [40]. The torsion scans are available on QCArchive https://qcarchive.molssi.org/ as a `TorsionDriveDataset` named `OpenFF Fragmenter Validation 1.0`.

### 5.4 Code and data availability

- Fragmenter code https://github.com/openforcefield/fragmenter
- Data generation, analysis scripts, and generating plots for the manuscript https://github.com/choderalab/fragmenter_data
- QCArchive dataset submission scripts https://github.com/openforcefield/qca-dataset-submission
- Exhaustive fragmentation validation https://doi.org/10.5281/zenodo.3987764

## Author Contributions

Conceptualization: CDS, CIB, JDC; Methodology: CDS, DGAS; Software: CDS, DGAS, LPW; Formal Analysis: CDS; Investigation: CDS; Resources: JDC; Data Curation: CDS, DGAS; Writing-Original draft: CDS; Writing-Review and editing: CDS, CIB, DGAS, JF, DLM, JDC; Visualization: CDS; Supervision: JDC, CIB, DLM, LPW; Project Administration: JDC; Funding Acquisition: CDS, JDC;

## Acknowledgments

We acknowledge Mehtap Isik, Andrea Rizzi, Gregory Ross, Bas Rustenberg, Patrick Grinaway, Melissa Boby, and other members of the Chodera lab for helpful discussions. We acknowledge Open Force Field Consortium members, specifically the scientists at Pfizer, Xinjun Hou, Brajesh Rai, and Qingyi Yang, and Caitlin Bannan, Alberta Gobbi, and Adrian Roitberg for helpful discussions. We acknowledge David Dotson for help in generating the fragmenter validation QC torsion scans.

## Disclaimers

The content is solely the responsibility of the authors and does not necessarily represent the official views of the National Institutes of Health.

## Appendix A Supplemental Figures

**Appendix A Figure 1.**
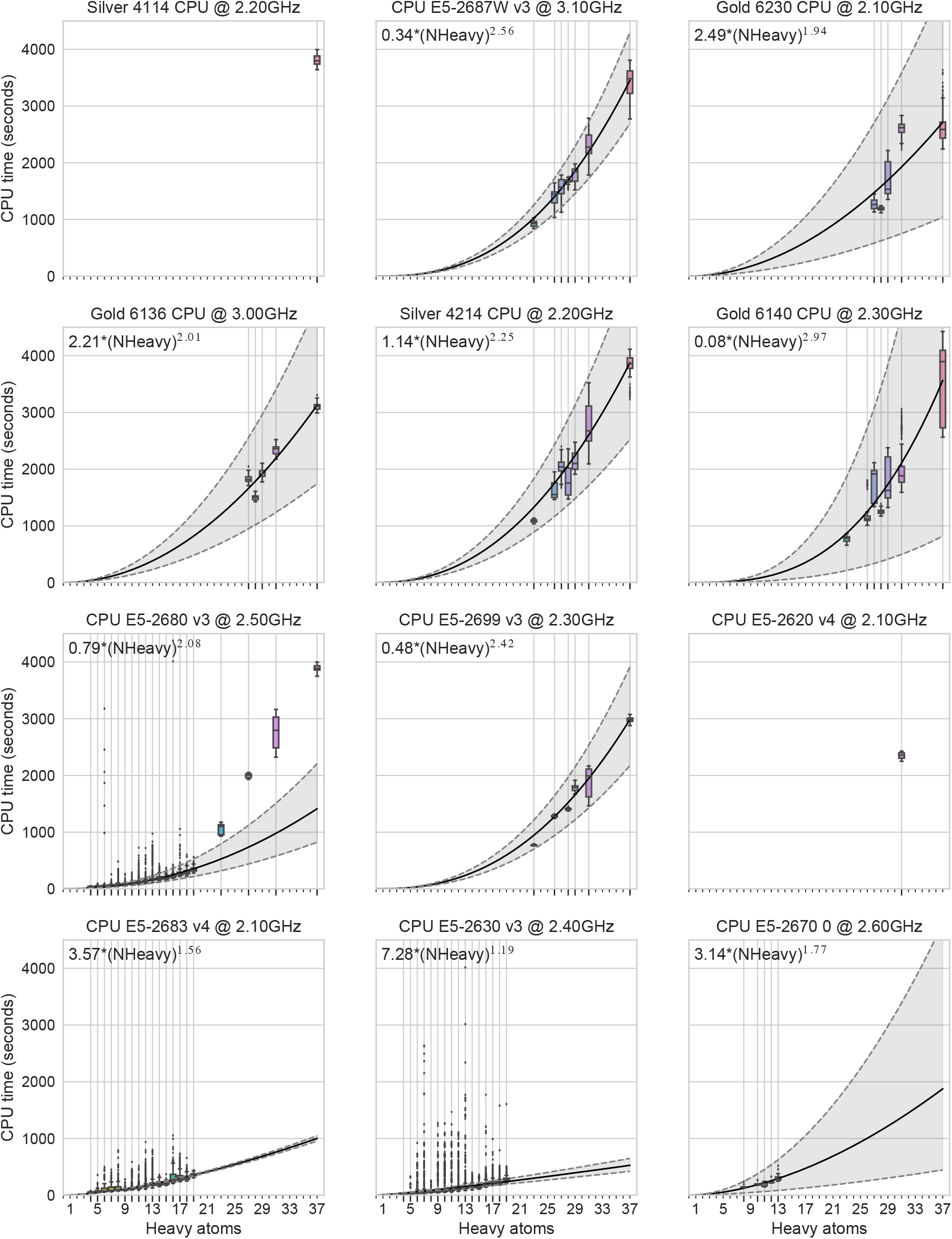
gradient evaluations scale similarly on various processor. CPU time (wall time * nthreads) for one gradient evaluation vs. heavy atoms in molecules. All CPUS shown in this figure are Intel(R) Xeon(R).

**Appendix A Figure 2.**
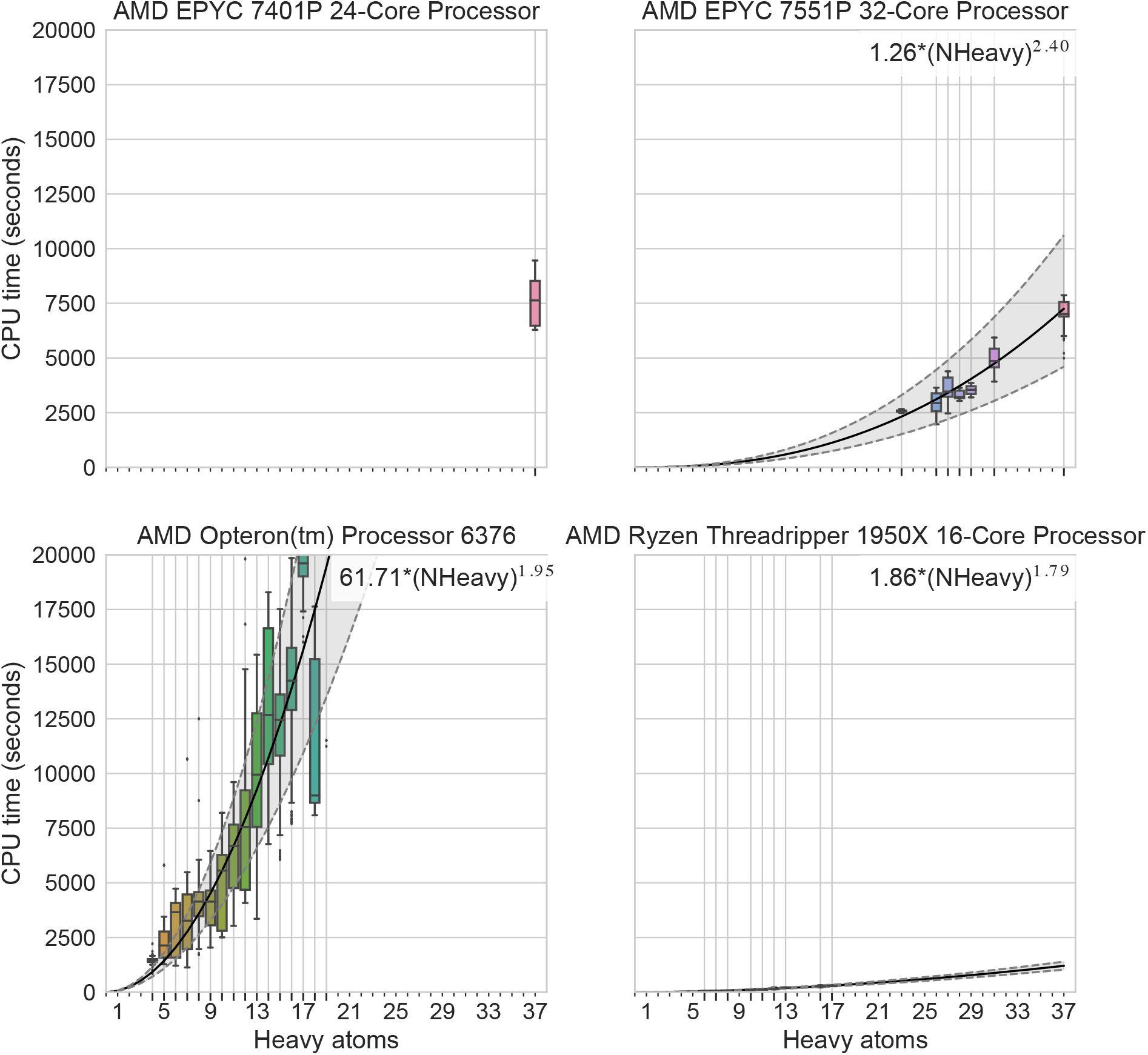
gradient evaluations scale similarly on various processor. CPU time (wall time * nthreads) for one gradient evaluation vs. heavy atoms in molecules. All CPUS shown in this figure are AMD processors.

**Figure 3.**
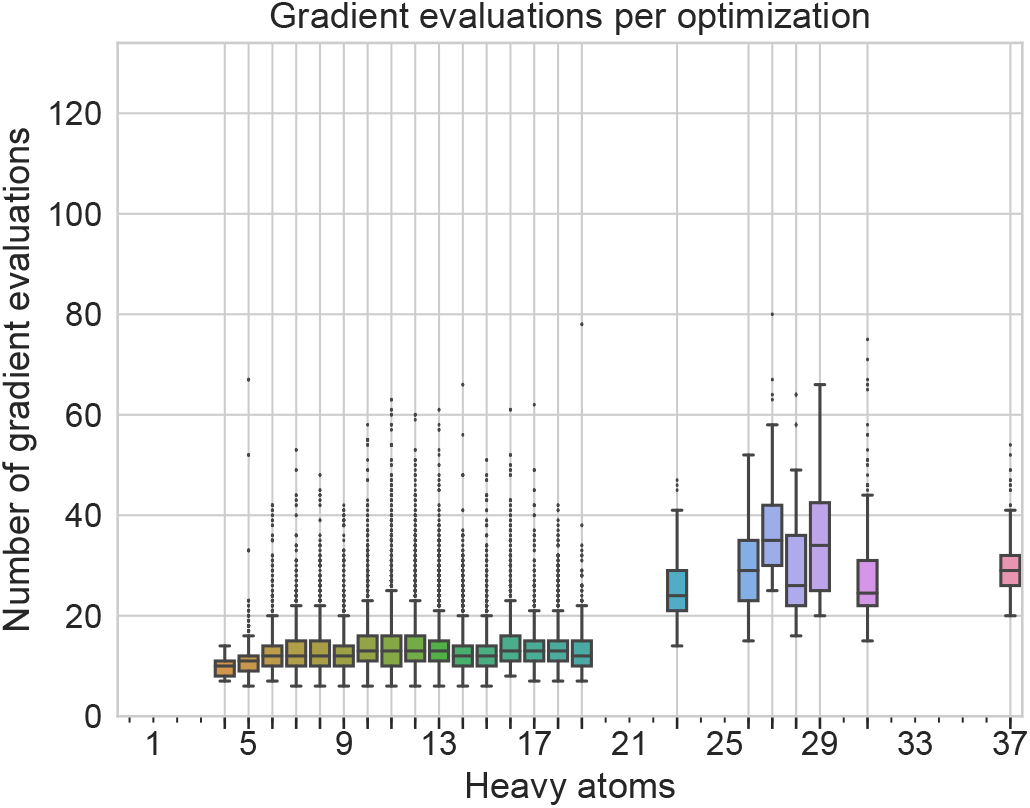
Distributions of number of gradient evaluations per optimizations for different size molecules. The number of gradient evaluations per optimization depends on many factors such as initialization and tolerance, but there is also a slight dependency on molecular size as shown in this figure.

**Appendix A Figure 4.**
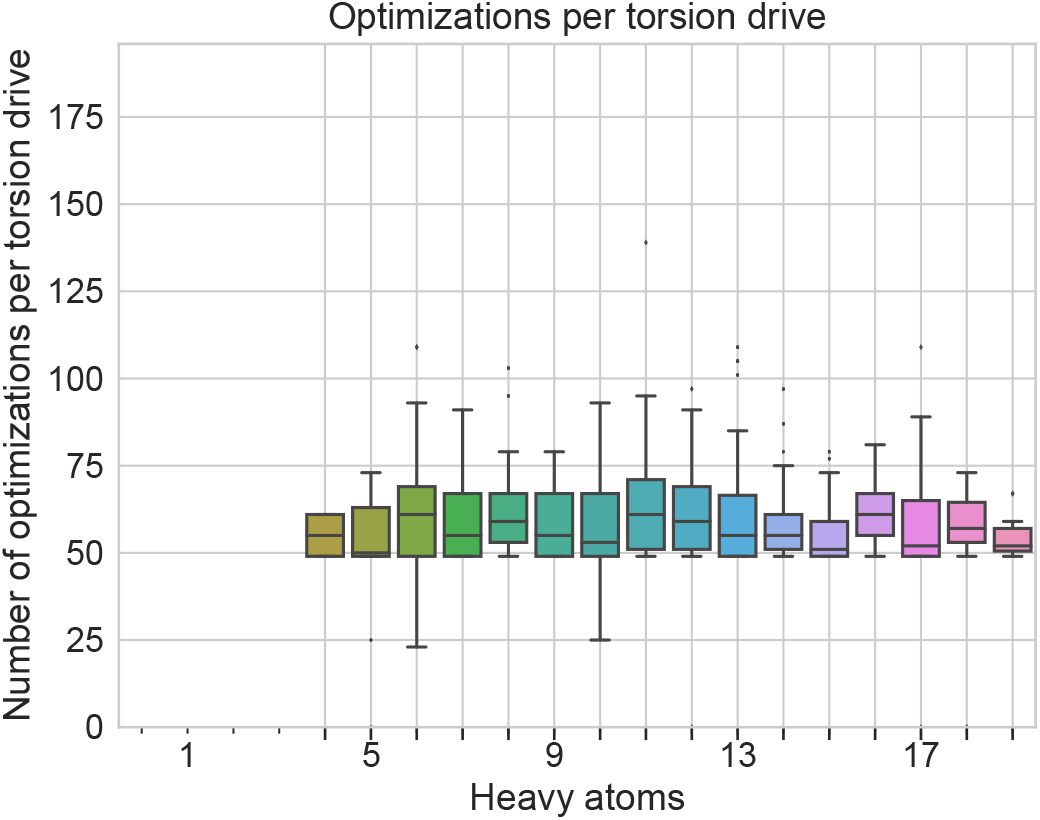
Distribution of optimizations per torsion drive. This figures shows the distributions of optimizations per torsion drive when using wavefront propagation.

**Appendix A Figure 5.**
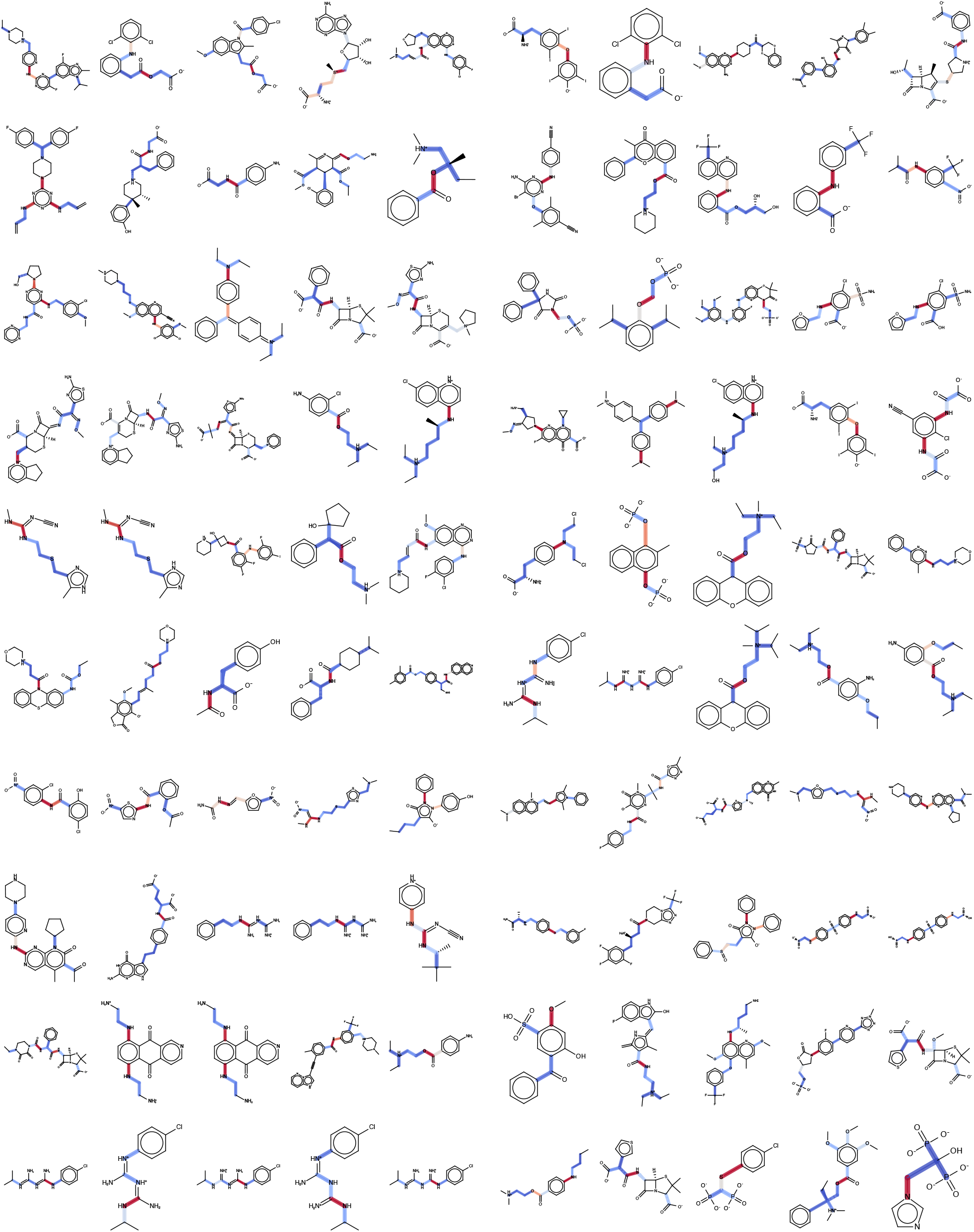
All molecules used in the validation set of fragmentation schemes. The bonds are highlighted by how sensitive they are to remote fragmentation. The redder bonds are more sensitive while the WBO distributions around the blue bonds do not change much with remote fragmentation.

**Appendix A Figure 6.**
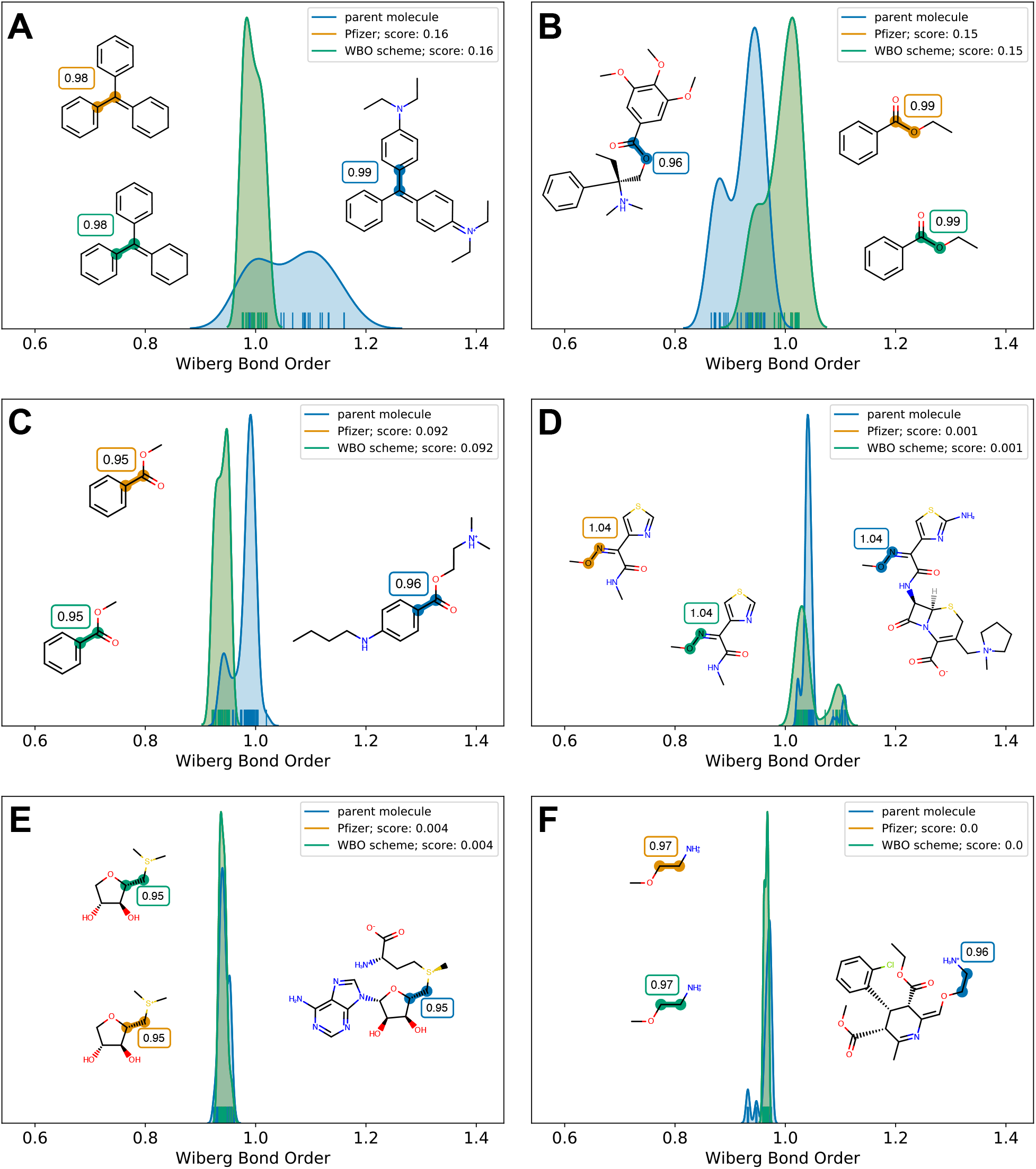
WBO fragmentation generates minimal fragment. This figure illustrates some cases where the Pfizer scheme and WBO scheme generate the same minimal fragment. While in most cases these fragments have WBO distributions that are close to the parent WBO distributions of the highlighted bond, sometimes both minimal fragments are equally poor. **[A], [B]**, and **[C]** show fragments of both schemes where important remote, chemical groups are not in the fragment so their overlapping distributions (orange and green) are far from the parent WBO distributions (blue distributions). **[D], [E]**, and **[F]** are the same as **A, B** and **C**, but the fragments do have the important chemical substituents so both perform equally well.

**Figure 7.**
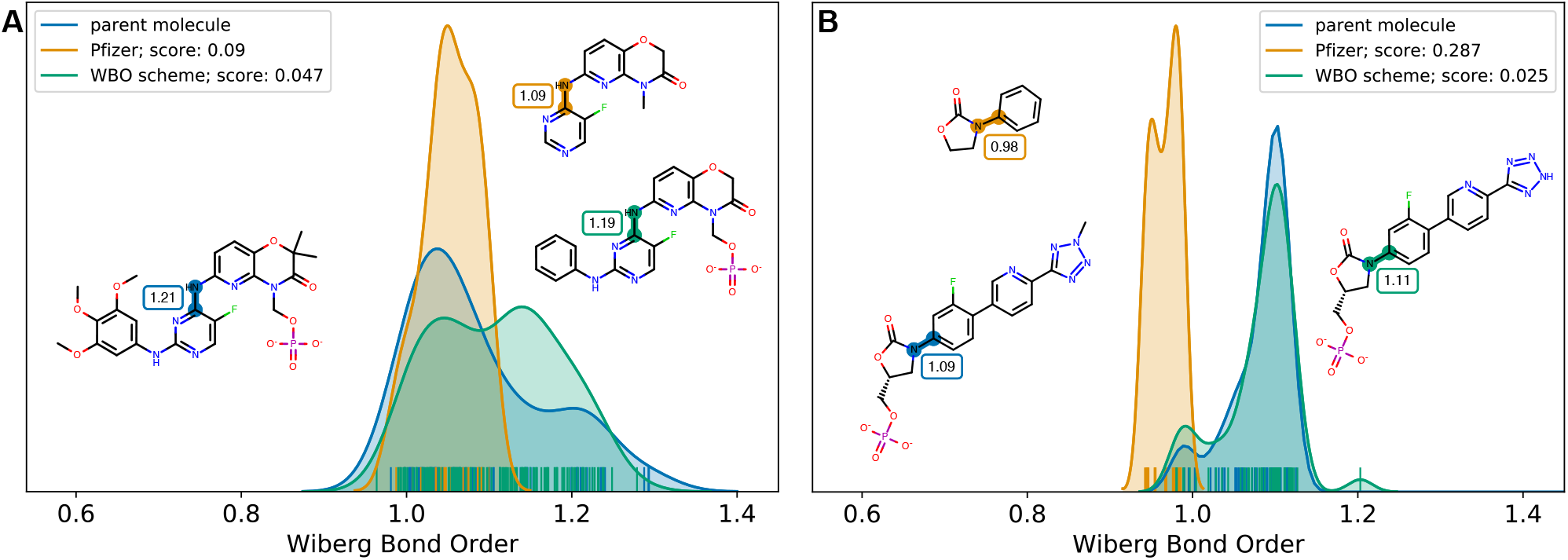
When Phosphate is removed, WBO distribution shifts even when the phosphate is six bonds away. Examples from benchmark set where remote phophsate groups induce long range effects of **[A]** six bonds away and **[B]** four bonds away.

**Appendix A Figure 8.**
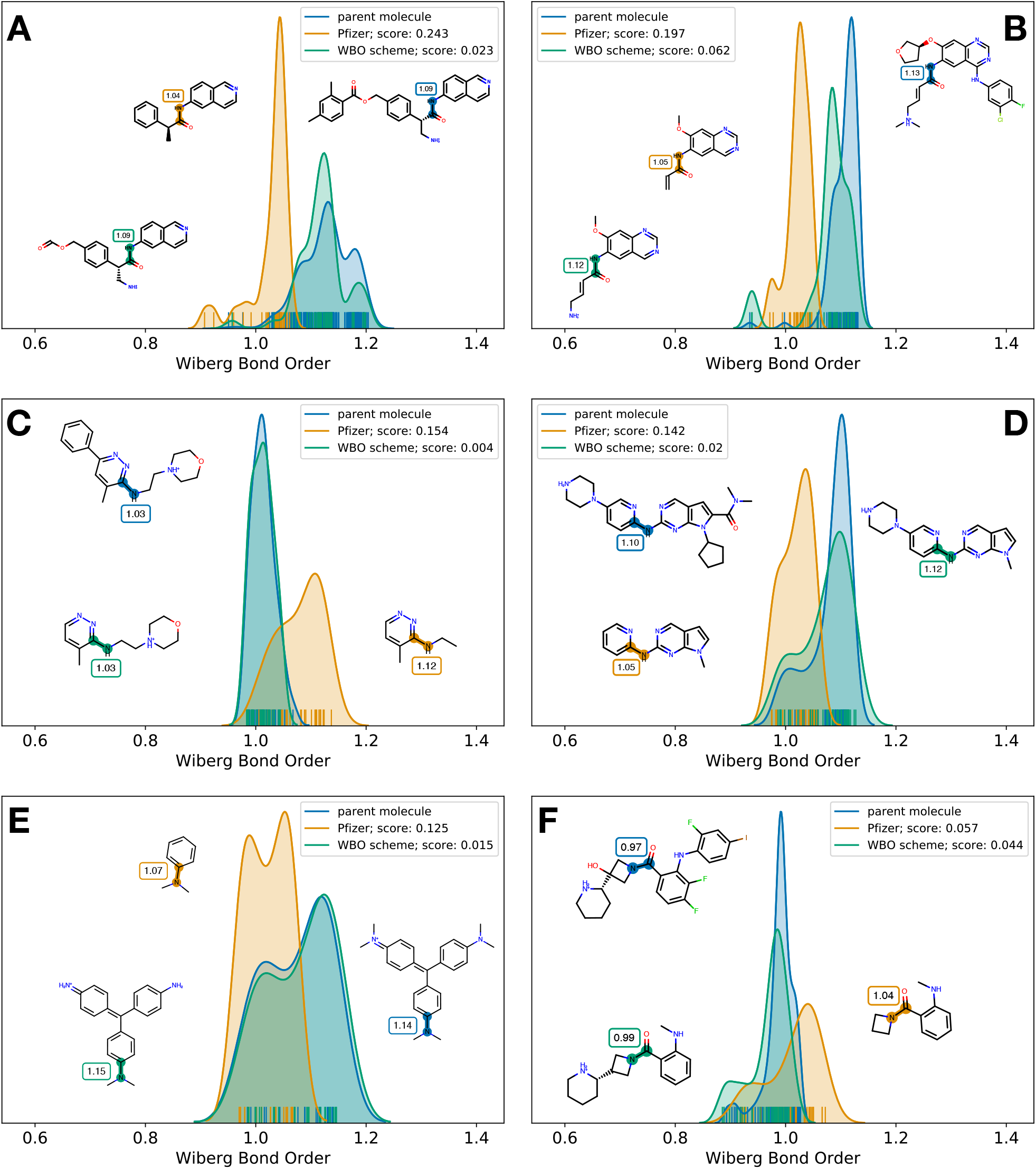
When a protonated nitrogen is removed, WBO distribution shifts even when the nitrogen is up to eight bonds away. Selected examples from the benchmark set were remote nitrogen groups induce long range effects. The distance of the remote nitrogen ranges from three bonds away up to eight bonds away.

**Appendix A Figure 9.**
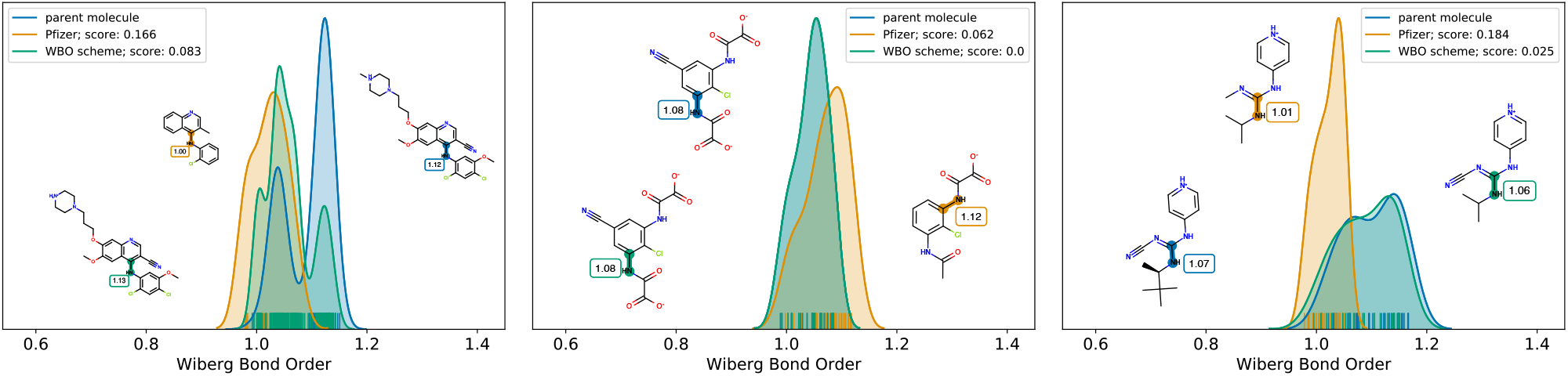
Removing a nitrile group shifts the WBO distribution. Examples from the benchmark set. Removing remote nitrile groups shift the WBO distribution.

**Appendix A Figure 10.**
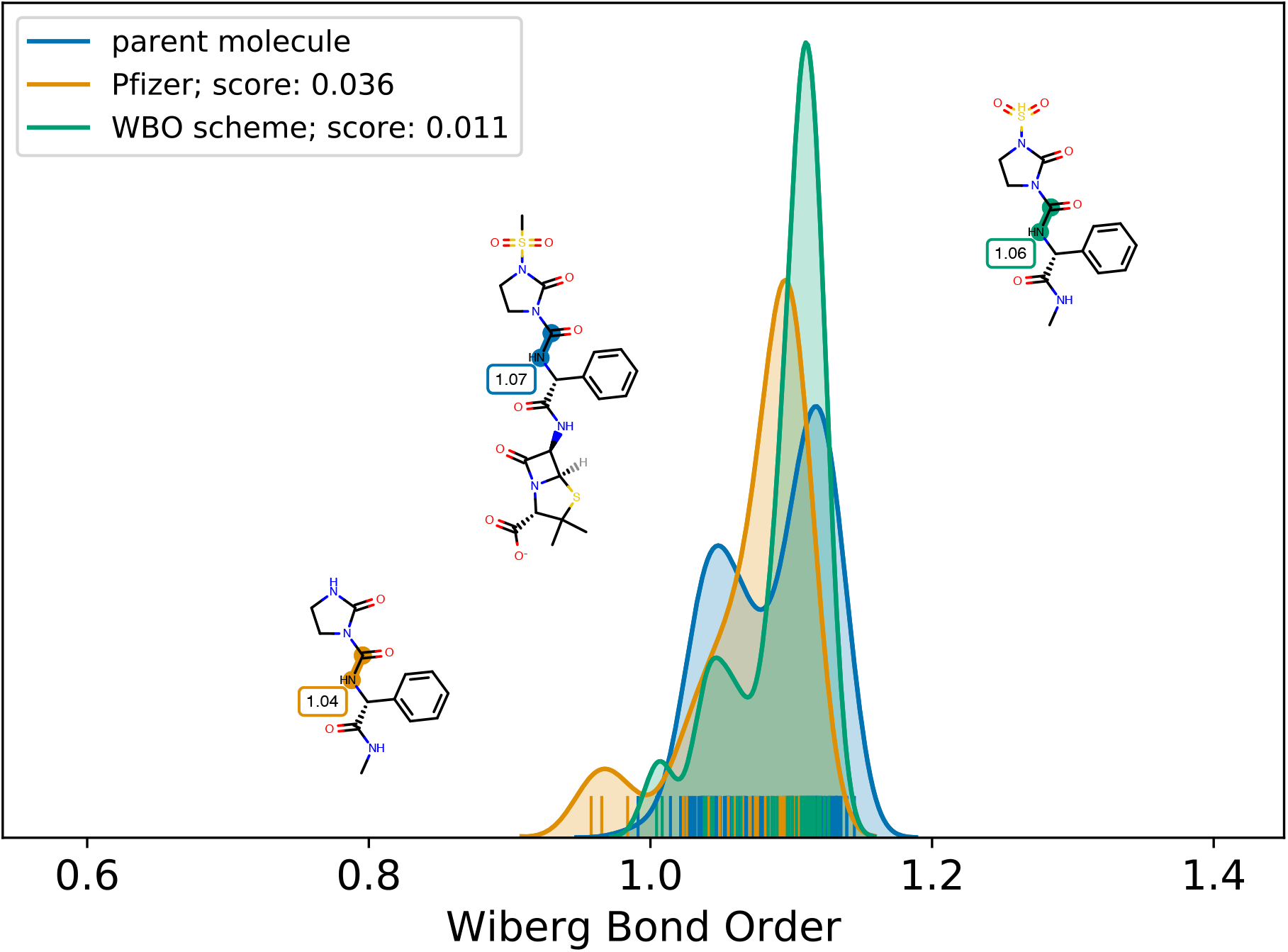
Removing a sulfonamide group shifts WBO distribution. Removing the sulfonamide (orange) that is four bonds away moves the fragment’s WBO distribution further away from parent WBO distribution (blue).

**Appendix A Figure 11.**
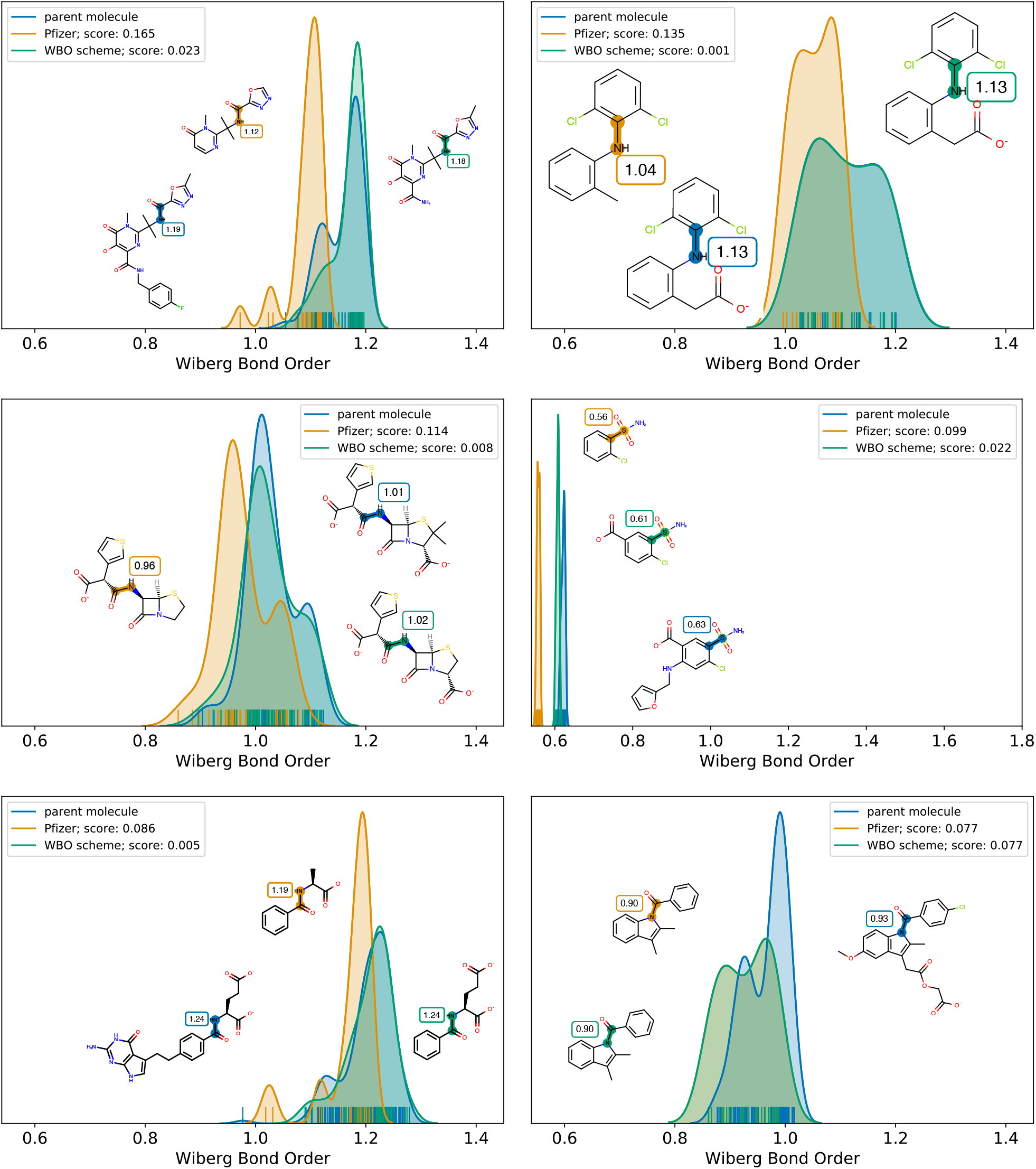
Removing a deprotonated oxygen shifts the WBO distribution. Removing a negatively charged nitrogen can effect bonds up to six to eight bonds away.

**Appendix A Figure 12.**
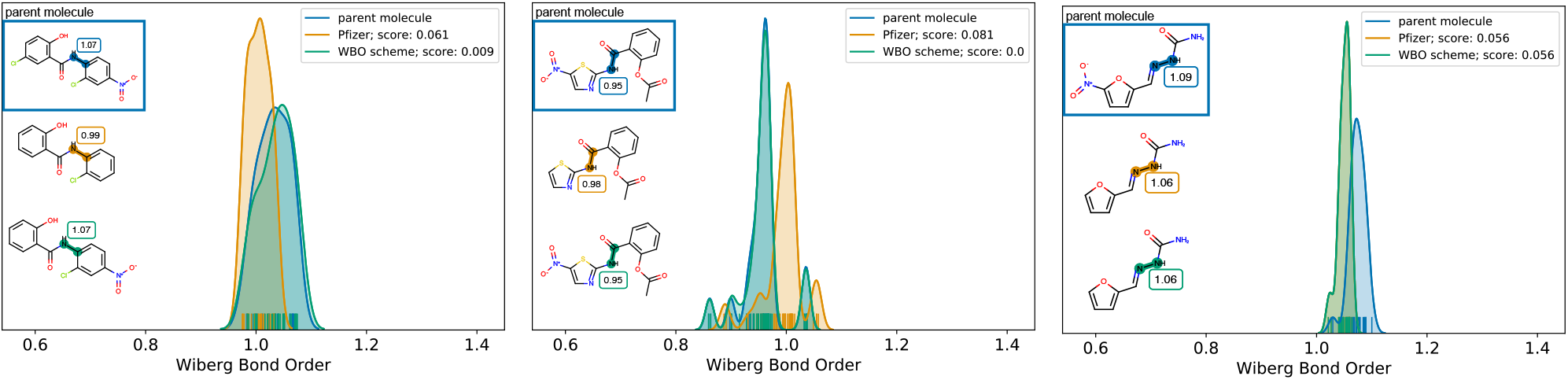
Removing a ntiro group shifts the WBO distribution. Removing a ntiro group effects bonds up to five bonds away.

**Figure 13.**
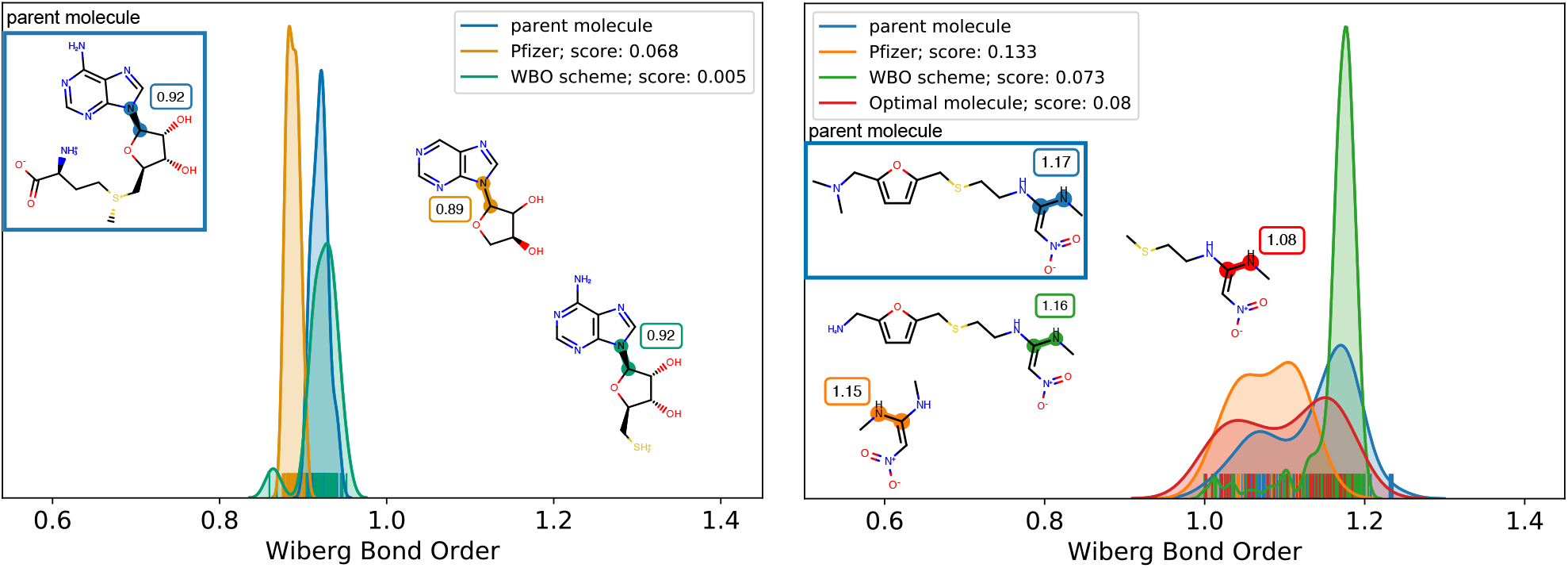
Removing sulfur shifts the WBO distribution. Removing sulfur shifts the WBO, but the shift is smaller than for other functional groups and this is not always captured by the WBO scheme. The figure on the right shows the fragment that the WBO scheme generates (green), however, it also includes the ring. The smaller fragment that has the sulfur (red) has an ELF10 WBO that is 0.1 lower than in the parent, but the WBO distribution (shown in red) is closer to the parent WBO distribution (blue, distance score 0.08) than the fragment without sulfur (orange, 0.133)

**Appendix A Figure 14.**
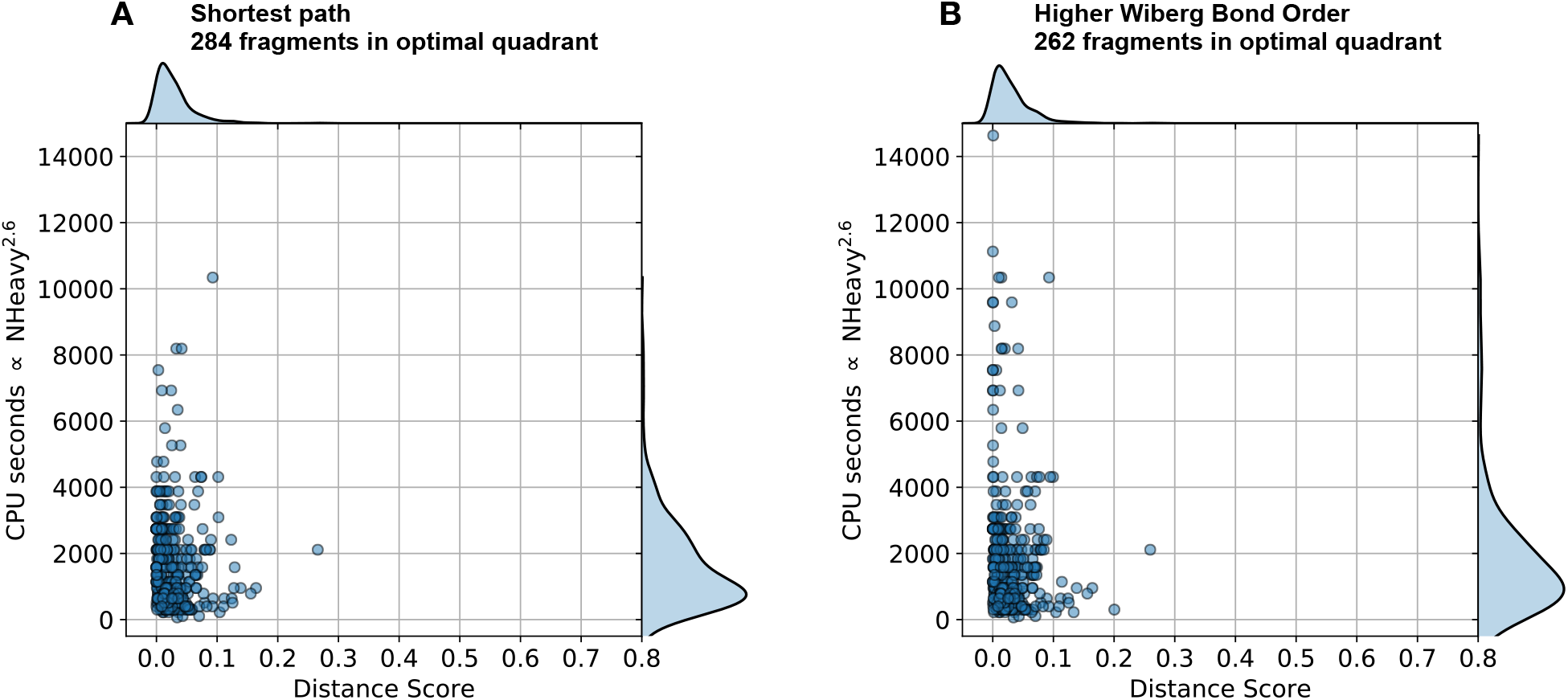
Comparing heuristics for growing out fragment. **[A]** Distance score and computational cost of fragments generated with the shortest path heuristic. **[B]** Same as **A** with the greater WBO heuristic.

## Footnotes

1 When performing discrete sampling of the torsional potential, it is desirable to sample near the critical points in order to obtain the qualitatively correct picture of the continuous potential energy curve. For 2-fold and 4-fold periodic potentials, we expect critical points at 0, 45, and 90 degrees, whereas for 3-fold and 6-fold periodic potentials, critical points are expected at 0, 30, 60, and/or 90 degrees. Therefore, the 15 degree grid spacing was chosen because it included most points where extrema in the torsional potential are likely to appear.

